# Endothelial Cells retain inflammatory memory through chromatin remodeling

**DOI:** 10.1101/2025.06.12.659136

**Authors:** Daniel Gonsales Spindola, Samantha Clark, Amber Bahr, Gabriel Pin de Jesus, Nina Martino, Antony Lowery, Shuhan Lyu, Andrew Seeman, Grace Martino, Giesse Albeche Duarte, Elijah Crosbourne, Peter Vincent, Guangchun Bai, Katherine MacNamara, Alejandro Adam, Ramon Bossardi Ramos

**Affiliations:** Albany Medical College; University at Buffalo – SUNY; Binghamton University

## Abstract

Sepsis survivors face a heightened risk of secondary infections and chronic vascular dysfunction despite clinical recovery, yet the underlying mechanisms remain poorly defined. Our study identifies a novel mechanism of endothelial inflammatory memory, wherein transient inflammatory exposure induces durable chromatin remodeling, priming endothelial cells for exaggerated responses to subsequent inflammatory insults. Utilizing a clinically relevant two-hit mouse model, cecal ligation and puncture (CLP) followed by secondary *Streptococcus pneumoniae* (SP) infection, we reveal persistent transcriptional activation in endothelial cells (ECs), characterized by amplified expression of inflammatory cytokines, adhesion molecules (e.g., ICAM-1), complement factors, and interferon-stimulated genes. Genome-wide ATAC-seq analyses demonstrated stable chromatin accessibility at key inflammatory loci, indicative of epigenetic priming. Mechanistically, we uncovered a critical role for the AP-1 transcription factor JunB in mediating this epigenetic remodeling. JunB knockdown attenuated chromatin accessibility and subsequent transcriptional amplification upon secondary challenge, pinpointing JunB-driven chromatin modifications as central to endothelial reprogramming. Our findings offer mechanistic insights into how transient inflammation creates lasting epigenetic states within the endothelium, highlighting JunB as a potential therapeutic target to mitigate chronic endothelial dysfunction and increased susceptibility to secondary infections post-sepsis.

## Introduction

Sepsis remains a major cause of morbidity and mortality globally, with survivors experiencing a significantly elevated risk of death from secondary infections (Shankar-Hari, Ambler et al. 2016). Although advances in critical care medicine have greatly improved survival rates during the initial acute hyperinflammatory phase of sepsis, a growing proportion of patients now survive only to face long-term sequelae and an increased vulnerability to secondary infections, particularly pneumonia (Wang, Derhovanessian et al. 2014, Shankar-Hari and Rubenfeld 2016, Zhao, Li et al. 2016, Pandolfi, Brun-Buisson et al. 2022, Inghammar, Linder et al. 2024). These secondary infections are closely associated with persistent endothelial dysfunction, characterized by impaired vascular integrity and unresolved inflammatory states, significantly contributing to prolonged organ dysfunction and increased late mortality (Ince, Mayeux et al. 2016, Fernandez, Palomo et al. 2021, Zhou, Liu et al. 2022). Importantly, current anti-inflammatory therapeutic strategies have proven ineffective in reducing the susceptibility to or severity of post-sepsis secondary infections(Kaneki 2017, Venet and Monneret 2018).

Endothelial dysfunction is central to the pathogenesis and prognosis of sepsis, where biomarkers of endothelial activation, such as interleukin-6 (IL-6) and intercellular adhesion molecule 1 (ICAM-1), correlate with worse clinical outcomes (Hotchkiss, Moldawer et al. 2016, He, Duan et al. 2024). Persistent endothelial activation exacerbates inflammatory responses, leading to increased vascular permeability, coagulopathy, leukocyte recruitment, and tissue hypoxia, which together perpetuate organ injury and dysfunction (Pober and Sessa 2007, Potter, Jiang et al. 2009, Joffre, Hellman et al. 2020, Augustin and Koh 2024).

While the critical role of endothelial cells (ECs) in acute inflammation is well established (Joffre, Hellman et al. 2020), their potential to retain a form of inflammatory memory that shapes responses to subsequent insults remains underexplored. Historically considered a feature of adaptive immune cells (Netea, Dominguez-Andres et al. 2020), it is now evident that innate and even non-immune cells— including ECs—can undergo durable functional reprogramming, a phenomenon referred to as “innate” or “trained” memory (Sohrabi, Lagache et al. 2020, Larsen, Cowley et al. 2021, Niec, Rudensky et al. 2021, Weiss, Vlahos et al. 2021, Lu, Sun et al. 2022, Naik and Fuchs 2022). In ECs, this memory is not mediated by antigen specificity but rather by persistent alterations in signaling cascades and transcriptional programs (Sohrabi, Lagache et al. 2020, Weiss, Vlahos et al. 2021, Lu, Sun et al. 2022). This maladaptive form of memory may underlie the heightened susceptibility to secondary infections and long-term organ damage observed in sepsis survivors, highlighting a critical yet largely unaddressed mechanism in post-sepsis pathology.

To investigate this hypothesis, we developed complementary in vitro and in vivo models of endothelial inflammatory memory. We employed an established mouse model of sepsis via cecal ligation and puncture (CLP), followed by a secondary mild intranasal challenge with *Streptococcus pneumoniae (SP)*, reflecting the clinical scenario of post-sepsis pneumonia. Performing CLP as a first hit, this model mimics the clinical response observed in sepsis patients promoting an acute inflammatory response in which a percentage of subjects will not survive, while the survivors exhibit a clinical recovery. In our model, the CLP survivors exhibit the recovery—including restored weight, temperature, and cytokine levels—they remain highly vulnerable to subsequent infection. Remarkably, secondary SP exposure triggered 84% mortality in CLP survivors, compared to 0% in sham controls, revealing a latent susceptibility. This was accompanied by an exaggerated systemic cytokine response and profound transcriptional reprogramming of endothelial cells in the lung and kidney. RNA-seq analyses demonstrated robust and organ-specific amplification of inflammatory, interferon-responsive, and endothelial activation pathways in CLP+SP mice, including elevated expression of ICAM1, C3, CCL2, and multiple interferon-stimulated genes. These amplified responses were paralleled by worsened renal function and increased ICAM-1 protein expression, establishing a mechanistic link between endothelial inflammatory memory and maladaptive host responses.

Concurrently, in vitro, human umbilical vein endothelial cells (HUVECs) were primed with IL-6 in the presence of receptor alpha (sIL-6Ra) to mimic a pro-inflammatory environment similar to the conditions observed in post-sepsis patients, followed by a rest period and a secondary inflammatory challenge with lipopolysaccharide (LPS) alone. Utilizing transcriptomic and epigenomic profiling, we identified sustained chromatin accessibility at critical pro-inflammatory gene loci (e.g. ICAM1 and CCL2) following initial inflammatory exposure. These genomic regions exhibit persistently increased chromatin accessibility, indicative of long-term transcriptional priming. Crucially, these epigenetic changes were associated with persistent activation of key transcription factors, notably STAT3 and JunB, which are known regulators of chromatin remodeling and inflammatory gene expression. These molecular signatures correlated with an amplified transcriptional response upon subsequent inflammatory challenges, strongly supporting the concept of inflammatory memory within endothelial cells.

Mechanistically, we directly investigated the functional relevance of JunB in sustaining endothelial inflammatory memory. Using targeted JunB knockdown (siJunB) in our in vitro endothelial model, we observed significant decrease in sustained chromatin accessibility at critical inflammatory genes. Moreover, JunB depletion substantially diminished the amplified transcriptional responses upon secondary inflammatory challenge, highlighting JunB as a necessary component in the establishment of endothelial inflammatory memory. Understanding the mechanisms underlying endothelial inflammatory memory is critical, as it holds potential implications for therapeutic interventions aimed at mitigating long-term vascular dysfunction in sepsis survivors. Targeting the molecular pathways driving this memory could represent a novel strategy to reduce susceptibility to secondary infections and improve long-term outcomes following severe inflammatory events.

## Methods

Mice C57BL/6J (Male or Female) were purchased from Jackson (Jackson Laboratory, Bar Harbor, ME, Cat# 000664) mice used in this study were bred and maintained under specific pathogen-free conditions in the Animal Research Facility at Albany Medical College. All experiments were carried out with 7-to 12-week-old mice according to the guidelines of the AMC Animal Care and Use Committee (Protocols number: 24-03004 and 24-08002). All animal experiments were approved and conducted in accordance with the Albany Medical College IACUC guidelines for animal care and were performed in the Animal Research Facility at Albany Medical College. Mice were housed in specific pathogen-free rooms with 12-hour light/12-hour dark cycle and controlled temperature and humidity. Mice were kept in groups of 5 or fewer in Allentown cages with access to food and water ad libitum.

### CLP-induced polymicrobial sepsis model

The CLP-induced polymicrobial sepsis mouse model was performed as described in previous works (Nascimento, Viacava et al. 2021, Biswas, Bahr et al. 2024). Mice were shaved and cleared of residual hair using depilatory cream, anesthetized with 1%–3% isoflurane administration, and placed in a supine position. The abdomen was disinfected using betadine and 70% ethanol, the abdominal cavity was opened, the cecum was exposed, and the distal half of the cecum was tightly ligated with 6-0 silk suture and punctured once with a 21-gauge needle to extrude a small amount of cecal content. The cecum was then returned to the peritoneal cavity, the peritoneum was closed with 6-0 silk sutures, and the skin was sealed using surgical glue (WEBGLUE, Pivetal, material # 21295419). Sham surgery was performed with surgical exposure of the cecum, but without ligation or puncture of the cecum. All mice received 0.1mL buprenorphine (0.5mg/kg, subcutaneously [s.c.]) immediately after and 6-10hrs post-surgery, as well as every 24hrs up to day 3. Mice received 0.1mL cefotaxime (50 mg/kg in 100 mL, s.c.) beginning 30 min after surgery and thereafter once per day for 3 days. Survival, severity score, body weight, and body temperature were observed for up to 20 days; mice experiencing surgical complications were excluded from analysis. On Day 20, the secondary hit of *SP* (1.0 x 10⁷) was administered intranasally to a portion of mice, while the remainder of mice who were not given SP treatment received saline intranasally as control and four groups were stablished: CLP+SP, CLP+Saline, Sham+SP and Sham+Saline.

### Glomerular Filtration Rate Measurement

Glomerular filtration rate (GFR) was assessed using a non-invasive fluorescence-based method with a transdermal Mini GFR Monitor (MediBeacon, St. Louis, MO, USA) as previously describe(Scarfe, Schock-Kusch et al. 2018, Martino, Ramos et al. 2021). One to two days prior to the procedure, a small area on the dorsal surface of each mouse was shaved and cleared of residual hair using depilatory cream. At 3 days or 15 days after CLP surgery, and at 24 hours following the secondary intranasal challenge with *SP* or PBS, mice were lightly anesthetized with 1-3% isoflurane and placed on a warming pad maintained at 37°C. The GFR monitor was affixed over the shaved region using medical-grade adhesive and tape. After a 1–2 minute baseline recording, FITC-sinistrin (7 mg per 100 g body weight) was administered via retro-orbital injection in sterile saline. Mice were then returned to individual cages without access to food or water during the 90-minute recording period. The monitor was subsequently removed, and GFR values were computed using proprietary analysis software provided by MediBeacon.

### Mouse Inflammation Panels

Cytokine levels in blood were quantified using the GeniePlex Mouse Inflammation assay (MOAMPM022, AssayGenie, Dublin, Ireland), following the manufacturer’s protocol. A total of 15µl of plasma per sample was used for the assay in duplicate. Samples were analyzed on an FACSymphony A3 flow cytometer (BD Biosciences, Franklin Lakes, NJ, USA). Data were processed and cytokine concentrations calculated using FCAP Array Software Version 3.0 (BD Biosciences) by an investigator blinded to experimental conditions.

### Endothelial Enrichment

PBS containing 0.1% BSA was prepared and Collagenase–Dispase (CD) mix was formulated by dissolving 300 mg Collagenase Type I and 327 mg Dispase II were dissolved in PBS. Streptavidin– antibody Dynabeads were prepared by resuspending a 100 µL aliquot in 1.4 mL PBS + 0.1% BSA, washing three times on a magnetic separator, and then coating the beads: 20 µL each of purified anti-EpCAM and anti-CD45. 20 µL of biotinylated anti-CD31 was added to the remaining tubes. After overnight incubation on a rotator at 4°C and four subsequent washes, the beads were resuspended in 100 µL PBS + 0.1% BSA and stored at 4°C.

Mice were euthanized via pentobarbital injection, and lungs and kidneys were excised, rinsed with cold PBS to remove blood, and kept on ice. Tissues were minced finely with scissors in separate dishes and transferred to 50 mL tubes containing 5 mL of pre-warmed (37°C) CD mix per mouse, with residual fragments washed into the tubes. Samples were incubated at 37°C with gentle agitation for approximately 30 min (mixing every 5 min), then triturated using a syringe with an 14G cannula followed by a 20G cannula until a uniform cell suspension was achieved. The suspension was filtered through a 70 µm cell strainer (washed with 5 mL PBS + 0.1% BSA), centrifuged at 400 × g for 10 min at 4°C, and the pellet was resuspended in 1.5 mL cold PBS + 0.1% BSA; 100 µL of this suspension was reserved for total RNA extraction, mixed with 500 µL TRIzol, the remaining 1.4 mL underwent cell separation. For negative and positive selection, 100 µL of streptavidin beads conjugated with biotinylated EpCAM and CD45 antibodies was added to the cell suspension and incubated for 15 min at room temperature on a rotator, followed by magnetic separation for 3 min to collect the supernatant; this supernatant was then incubated with 100 µL beads conjugated to biotinylated CD31 for 15 min at room temperature, magnetically separated, and washed six times with 1 mL PBS + 0.1% BSA. The bead-bound cells were resuspended in 100 µL sterile PBS, transferred to RNase-free tubes, mixed with 500 µL TRIzol, and processed for RNA isolation.

RNA extraction for both total and bead-bound fractions involved thawing the TRIzol samples for 5 min at room temperature, mixing thoroughly, adding 100 µL chloroform. The upper aqueous phase (∼300 µL) was transferred to a new tube, combined with 300 µL of 70% ethanol, and loaded onto RNeasy MiniElute columns, with subsequent washes and RNA was eluted in 14 µL RNase-free water and quantified by NanoDrop, with the resulting RNA samples proceeding to RNA-seq analysis.

### HUVEC Two hit model treatment

HUVEC were isolated in-house according to established protocols as described (Martino, Ramos et al. 2021, Martino, Bossardi Ramos et al. 2022, Bossardi Ramos, Martino et al. 2023). To model inflammatory memory in human endothelial cells, HUVECs were cultured in 12-well plates at a density of 320,000 cells per well in EGM2 medium. Cells were stimulated with recombinant human IL-6 (200 ng/mL final) in combination with soluble IL-6 receptor (sIL-6R, 100 ng/mL final) for 72 hours to mimic the first inflammatory hit. After this initial stimulation period, cells were either harvested immediately or subjected to a 48-hour washout phase in cytokine-free medium to allow for resolution of acute signaling.

To simulate a second inflammatory challenge, a subset of washed cells was subsequently treated with lipopolysaccharide (LPS, 1 μg/mL) for 6 hours. Control groups included PBS-treated cells with and without LPS exposure. At each stage—post-IL-6 treatment, after the washout period, and following the LPS challenge—cells were collected, either for cryopreservation to ATAC-seq or RNA extraction. For ATACseq, cells were trypsinized, counted, and transferred to cryopreservation medium containing 10% DMSO before being stored at −80°C. Remaining cells were lysed in TRIzol reagent and stored at −80°C for RNA extraction following company protocol.

### RT-qPCR

Total RNA was isolated from either cultured HUVECs or mouse endothelial cells in single-cell suspension using TRIzol reagent (Thermo Fisher Scientific), following standard phenol-chloroform extraction and isopropanol precipitation protocols. For cDNA synthesis, 400 ng of total RNA was reverse transcribed using the PrimeScript RT Master Mix kit (Takara) at 42°C, as recommended by the manufacturer. The resulting cDNA was diluted 1:10 in nuclease-free water, and 2 μL of this diluted product was used per quantitative PCR reaction. Reactions were carried out using a StepOnePlus real-time PCR system (Applied Biosystems) with SYBR Green iTaq Universal Supermix (Bio-Rad), gene-specific primers (2 pmol), and nuclease-free water in a total volume of 20 μL. Relative gene expression levels were determined using the ΔΔCt method, normalized to GAPDH for human samples or HPRT for mouse samples.

### Gene Silencing via siRNA Transfection

For targeted knockdown of JunB, HUVECs were transfected with a pool of four siRNAs (ON-TARGETplus SMARTpool, Horizon Discovery) using Lipofectamine RNAiMAX (Invitrogen) in suspension. Non-targeting control siRNAs from the same vendor were used as negative controls. Prior to transfection, cells were lifted in 5% Opti-MEM (1% FBS in Opti-MEM) to avoid the presence of heparin from standard EGM2 medium, which could interfere with transfection efficiency. For each condition, siRNA and Lipofectamine were pre-incubated in Opti-MEM to form complexes at a final siRNA concentration of 150 nM. The complexes were allowed to form for 5 minutes at room temperature before being mixed 1:1 with the cell suspension and plated directly.

After 6 hours, EGM2 medium supplemented with penicillin/streptomycin and serum was added to each well to support cell recovery. Cells were maintained under standard culture conditions and subsequently treated with the two hit model treatment as described above. Total RNA was harvested in TRIzol, and protein lysates were prepared in Laemmli buffer. Knockdown efficiency of JunB was confirmed by quantitative RT-PCR and Western blotting.

### Western Blot

The gels were made using a casting kit (TGX Stain-Free FastCast Acrylamide Kit, 10%, BioRad, Cat. #1610183), 10% APS (Ammonium Persulfate, BioRad, cat. #1610700) and TEMED (TEMED, BioRad, cat. #161-0800). Once polymerized, the gels were loaded into BioRad Mini-PROTEAN Tetra Cell apparatus (cat. #1658005) and lysate samples that were stored in Laemmli buffer were pipetted 15uL per well. The gels ran for 90 minutes on 150volts before they were removed and transferred on top of a layer of filer paper and nitrocellulose membrane soaked for at least 2 minutes in Transfer Buffer (Trans-Blot Turbo RTA Mini Transfer Kit, BioRad, cat. #1704272), with another layer of filter paper on top, and then placed into BioRad machine for 10 minutes. Once the gel was transferred onto the membrane, it was stained using Ponceau (Boston BioProducts Inc, cat. #ST-180) and cut to size depending on the kDa banding of the preferred antibody. The membrane was then placed in 5mL of the proper blocker (either 5% BSA or Dry Milk) and placed on a rocker at room temperature for 1 hour. After an hour, the primary antibody was added at the proper dilution to the nitrocellulose, and it was then placed in a cold room to rock overnight. The next day, the membranes were washed 3 times for 10 minutes with PBS Tween (10X PBS, BioRad, ca. #1610780), before the secondary antibody was added at a 1:500 ratio and then rocker at ambient temperature for an hour. After the hour, the membrane rocked for 20 minutes in PBS Tween before being taken to the ChemiDoc to be read using equal parts of BioRad Clarity (Clarity Western ECL Substrate, BioRad, cat. # 170-5061). The following primary antibodies were used: anti-MCP1 (Cell Signaling Technology, cat. #81559T, 1:1000, blocked with dry milk), anti-β-ACTIN (Sigma-Aldrich, cat. #A5441, 1:5000, blocked with BSA), anti-STAT3 (Cell Signaling Technology, cat. #9145, 1:1000, BSA), anti-STAT3α (Cell Signaling Technology, cat. #8768, 1:1000, BSA), and anti-TNFAIP3 (Cell Signaling Technology, cat. #5630S, 1:1000, BSA).

### RNA-Seq

The bulk RNA-seq data discussed in this publication have been deposited in NCBI’s Gene Expression Omnibus and are accessible through GEO Series accession number GSE298859 for the 2-hit model on HUVEC, GSE299114 for the lung and kidney ECs, and GSE299261 for the siRNAseq for the JunB knockout HUVEC. Bulk RNA-seq data from endothelial cells subjected to a two-hit inflammatory model were processed and analyzed using a standardized pipeline in R. Total RNA was extracted using TRIzol and purified with the RNeasy Mini-Elute kit (Qiagen), followed by quality and concentration assessment with a NanoDrop spectrophotometer. RNA-seq libraries were prepared using Illumina TruSeq protocols and sequenced on an Illumina NextSeq 500 platform. Raw reads were aligned to the mm10 (mice) or hg38 (human) reference genome using the align() function from the Rsubread package (v2.10.5) with default parameters(Liao, Smyth et al. 2019). Gene-level counts were quantified using featureCounts, using the inbuilt gene annotation based on Entrez Gene IDs. Genes were filtered to retain those with a count-per-million (CPM) > 0.5 in at least 3 samples. Gene symbols were assigned using NCBI annotations.

Filtered count matrices were transformed into log2-CPM values using the edgeR package and normalized using the trimmed mean of M-values (TMM) method (Chen, Chen et al. 2025). Differential expression analysis was conducted with the limma-voom pipeline, incorporating empirical Bayes moderation to assess statistical significance (Law, Chen et al. 2014). Genes with an adjusted p-value < 0.05 and |log2 fold change| > 0.5 were considered significantly differentially expressed. Principal component analysis (PCA) was performed using base R and prcomp() to assess global transcriptional variation.

Gene set enrichment analysis (GSEA) was performed using WebGestalt (Elizarraras, Liao et al. 2024) to assess biological pathways significantly enriched in differentially expressed genes across experimental conditions. Enrichment was computed using the GSEA method with the Gene Ontology databases (Biological Process) and Pathway (KEGG) as reference sets. Adjusted p-values were calculated and categories with FDR < 0.05 were considered significant. Only genes on the amplified response with adjusted p-value < 0.05 were included as input. To infer transcription factor (TF) activity from bulk RNA-seq data, we implemented the DoRothEA framework (Garcia-Alonso, Holland et al. 2019, Badia, Velez Santiago et al. 2022). A confidence-weighted regulon (levels A–C) for human or mouse was applied depending on the organism, and normalized gene expression matrices were used to estimate enrichment scores for each TF. Enrichment scores (NES) > 2 were considered indicative of TF activation. This analysis enabled identification of key regulators underlying differential transcriptional states.

### ATAC-seq

The ATAC-seq data discussed in this publication have been deposited in NCBI’s Gene Expression Omnibus and are accessible through GEO Series accession number GSE298902 for the 2-hit model on HUVEC. ATAC-seq was performed at the Cornell University Biotechnology Resource Center (BRC) Epigenomics Facility (RRID:SCR_021287). Briefly, 100k cells flash-frozen in 10% DMSO were lysed, permeabilized, and tagmented using the Omni ATAC-seq protocol (Grandi, Modi et al. 2022), followed by 12 cycles of barcoding PCR. Gel-purified libraries were sequenced on the Element Biosciences AVITI to obtain at least 10M paired-end reads. Raw BAM files were initially processed by removing mitochondrial reads (chrM) using samtools and blacklisted genomic regions using bedtools intersect (Li, Handsaker et al. 2009, Quinlan and Hall 2010). Filtered BAM files were used for peak calling with MACS3 (callpeak), specifying parameters for paired-end data, genome size, and adjusted p-value threshold (q-value <0.1) (Zhang, Liu et al. 2008). Peaks were further assessed for quality by computing the fraction of reads in peaks (FRiP score). For data visualization, BAM files were sorted and indexed (samtools sort and samtools index), followed by the generation of BigWig coverage tracks normalized to counts-per-million (CPM) using bamCoverage (Yan, Powell et al. 2020).

Consensus peak sets were generated by merging peaks across replicates and conditions using bedtools merge, initially creating consensus peaks per experimental condition, and subsequently producing a unified peak set encompassing all experimental groups. Peak occupancy counts per sample were quantified using bedtools multicov, resulting in a final peak-count matrix utilized for downstream analyses of chromatin accessibility(Landt, Marinov et al. 2012).

For the elbow plot, we first generated a master set of peaks by merging all consensus peaks from both conditions. For each peak, we used DESeq2 to estimate the log2 fold change (log2FC) of accessibility in IL6 72h versus PBS 72h. We then computed a score by dividing the log2FC by its standard error (SEM), effectively producing the Wald statistic for each peak. To visualize these results, we ranked all peaks from highest to lowest based on their log2FC/SEM score and plotted them along the x-axis. The y-axis represents the log2FC/SEM value itself. Peaks with positive log2FC/SEM scores (above a chosen threshold) were classified as “open IL6 72hrs,” while those with negative scores (below a chosen negative threshold) were classified as “close IL6 72h.” Peaks whose scores remained near zero were considered “no changes.” Horizontal lines were drawn at the thresholds to delineate these categories.

Peaks were imported and merged within each condition using genomic-range operations (Bioconductor package GenomicRanges). Consensus peak sets were established by merging overlapping peaks from replicates. Peaks uniquely present in IL-6-treated samples (72h), absent in PBS controls, were identified using set-difference operations (setdiff) (Lawrence, Huber et al. 2013). To determine persistence, these IL-6-specific peaks were intersected (subsetByOverlaps) with peaks detected after the washout period (IL-6 72h + wash). Overlapping regions were defined as peaks “remaining open.” Peak overlaps and set relations were visualized using Venn diagrams (VennDiagram) (Chen and Boutros 2011). Persistent peaks were exported in BED format and annotated using the ChIPseeker package, mapping peaks relative to transcription start sites (TSS, ±2kb) based on UCSC annotations (Wang, Li et al. 2022). Annotated peaks were mapped to gene symbols using Ensembl database annotations. Multiple peaks assigned to the same gene were consolidated into single genomic regions spanning from the earliest start to the latest end positions.

The resultant gene set was integrated with RNA-seq differential expression results to correlate persistent chromatin accessibility with transcriptional changes. Specifically, genes exhibiting persistent chromatin opening after washout were merged with genes significantly upregulated in RNA-seq analysis following the two-hit inflammatory challenge.

## Results

### Two hit model of infection

To model the long-term consequences of sepsis, we employed a two-hit murine system beginning with surgical cecal ligation and puncture (CLP), a well-established model of polymicrobial sepsis or a sham surgery as control. Mice receive analgesics and antibiotics on days 1–3 to mimic clinical management. Glomerular filtration rate (GFR) and plasma are assessed on days 13 and 15, respectively, to confirm resolution of acute illness. On day 20, all mice are intranasally challenged with either *SP* or saline as a second hit. Final GFR and plasma samples are collected on days 21 (one day after *SP)* and 22 (two days after *SP*) to evaluate physiological responses to the secondary challenge (**Fig. 1A**).

**Fig 1.**
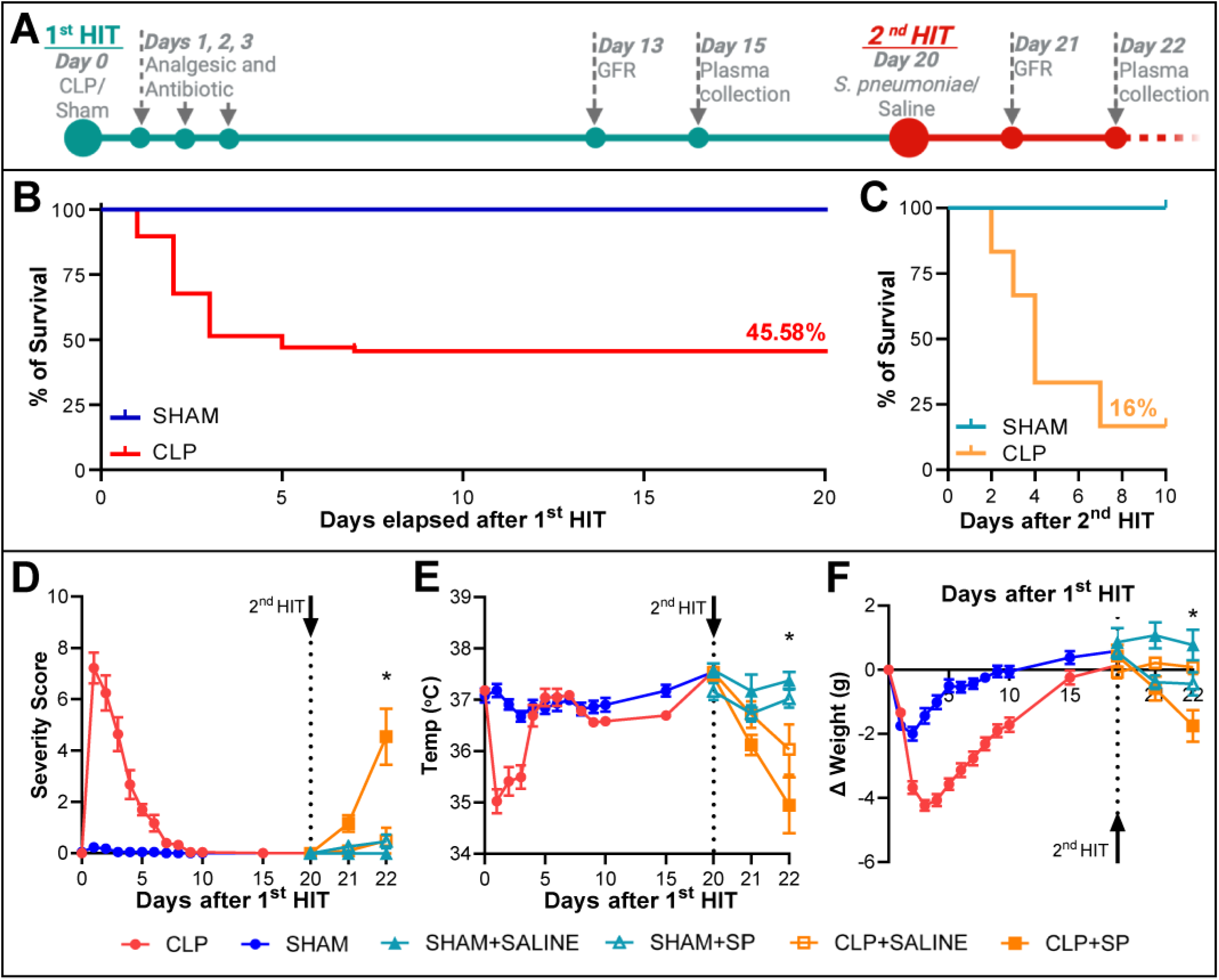
Two-hit Sepsis Model Induces Persistent Vulnerability and Exacerbated Secondary Response. **(A)** Experimental timeline: mice underwent cecal ligation and puncture (CLP) or sham surgery (first hit), mice that recovered clinically by day 20 was performed secondary intranasal challenge with S. pneumoniae (*SP*). **(B)** Survival curve post-CLP shows 45.6% mortality by day 8. **(C)** Survival following *SP* challenge reveals 16% mortality exclusively in CLP survivors. **(D)** Severity score, temperature, and weight changes confirm worsened clinical condition specifically in the CLP+SP group after secondary hit. p<0.05 by log-rank test (survival curves) or two-way ANOVA with post hoc comparisons.

As expected, CLP (1^st^ hit) induced acute illness with 54.4% survival, with most mortality occurring between days 3 and 8 (**Fig. 1B**). However, despite appearing clinically recovered, these sepsis survivors demonstrated profound susceptibility when re-challenged with an intranasal dose of *SP* on day 20. While this secondary infection was non-lethal in sham-operated controls, it induced 84% mortality in the CLP survivor group within four days (**Fig. 1C**), revealing a hidden vulnerability.

Mice that survived this initial insult showed clinical recovery by multiple physiological metrics, including normalized severity scores, temperature, and body weight by day 15 post-CLP (**Fig. 1D/E/F**). Notably, upon second hit, CLP+*SP* mice exhibited a dramatically worsened physiological response compared to SHAM+*SP* animals, with significant deterioration in clinical scores, temperature regulation, and weight loss (**Fig. 1D/E/F**).

Further, we quantified plasma cytokines at three critical time points: day 3 (acute sepsis phase), day 15 (clinical resolution phase), and day 22 (two days post-secondary *SP* challenge). During acute sepsis, we observed a significant elevation of pro-inflammatory cytokines in the CLP group, including IL-6, CCL2 (MCP-1), CXCL1 (KC), and CXCL10 (IP-10), indicating a strong systemic inflammatory response. By day 15, cytokine levels returned to baseline and were no longer significantly different from sham controls (p > 0.05), consistent with apparent clinical recovery. However, following the secondary *SP* challenge, CLP survivors exhibited a markedly amplified cytokine response. Plasma concentrations of IL-6, CCL2, CXCL1, and CXCL10 surged well beyond both resolved and acute-phase levels, revealing a reactivation of the inflammatory program uniquely triggered in post-septic mice upon re-exposure to infection (**Fig 2**).

**Fig 2.**
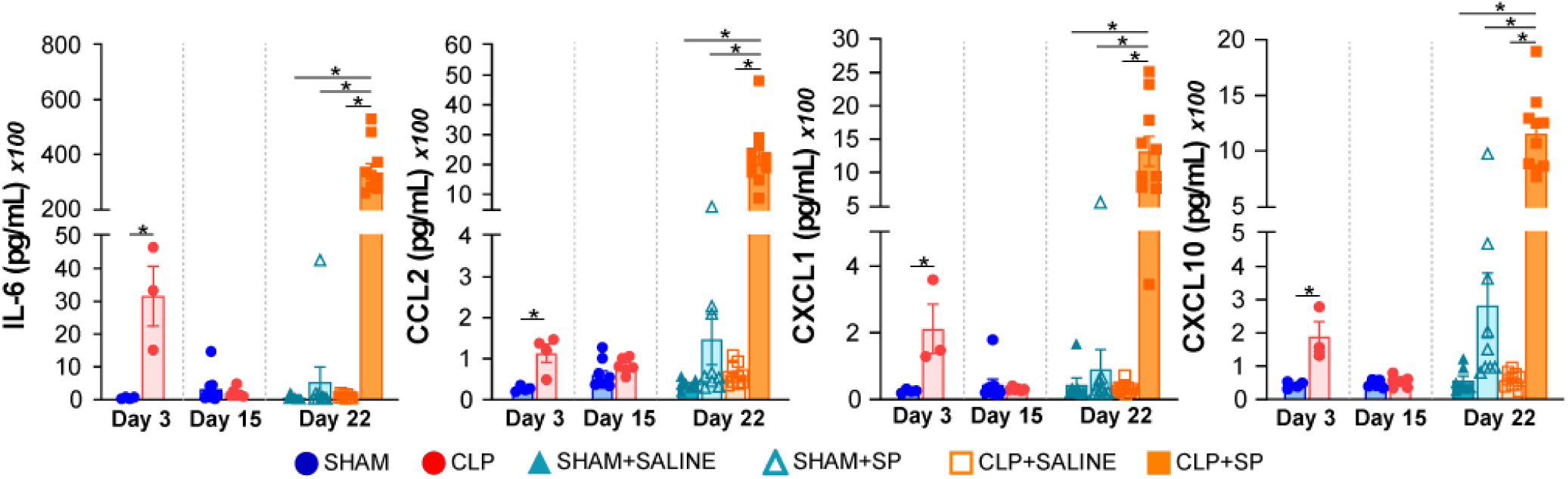
Amplified Systemic Cytokine Response upon Secondary Infection. Plasma cytokine levels (IL-6, CCL2, CXCL1, CXCL10) measured during acute sepsis (day 3), clinical recovery (day 15), and two days post-secondary SP challenge (day 22). Marked cytokine amplification is specifically observed in CLP survivors after secondary infection, significantly surpassing acute-phase levels. Data shown as mean±SEM; *p<0.05 by two-way ANOVA and post hoc analysis.

These data indicate that while CLP survivors meet standard criteria for recovery, they remain functionally compromised and primed for severe outcomes upon a secondary infectious insult. This model recapitulates the clinical scenario of post-sepsis immunopathology and provides a robust platform to explore the mechanistic underpinnings of inflammatory memory and maladaptive responses in endothelial and immune compartments.

### Lung Endothelial Transcriptional Activation Following Secondary Pulmonary Challenge

Given that our secondary insult is an intranasal challenge with *SP*, we first evaluated lung endothelial transcriptomic responses as the primary infection site. To assess endothelial-specific transcriptional responses, we isolated RNA from magnetically enriched ECs from lungs on day 3 and 22 after CLP, following our previously validated protocol (Bossardi Ramos, Martino et al. 2023, Lu, John Portela et al. 2024). Enrichment efficiency was confirmed by qPCR, showing robust depletion of the epithelial marker Cdh1 and significant enrichment of the endothelial marker von Willebrand factor (vWF) in the post-enrichment samples compared to pre-enrichment input (**Supplemental Fig 1**). This validated the specificity of our EC population and ensured that downstream transcriptomic profiling reflects endothelial-intrinsic gene expression changes.

Principal component analysis (PCA) revealed distinct transcriptional signatures at each phase of the experimental model, clearly separating acute CLP (day 3) from recovered (day 20) and second-hit CLP+SP conditions (**Fig. 3A**). The CLP+SP endothelial cells exhibited the most pronounced transcriptional changes, highlighting a unique response pattern upon secondary infection.

**Fig 3.**
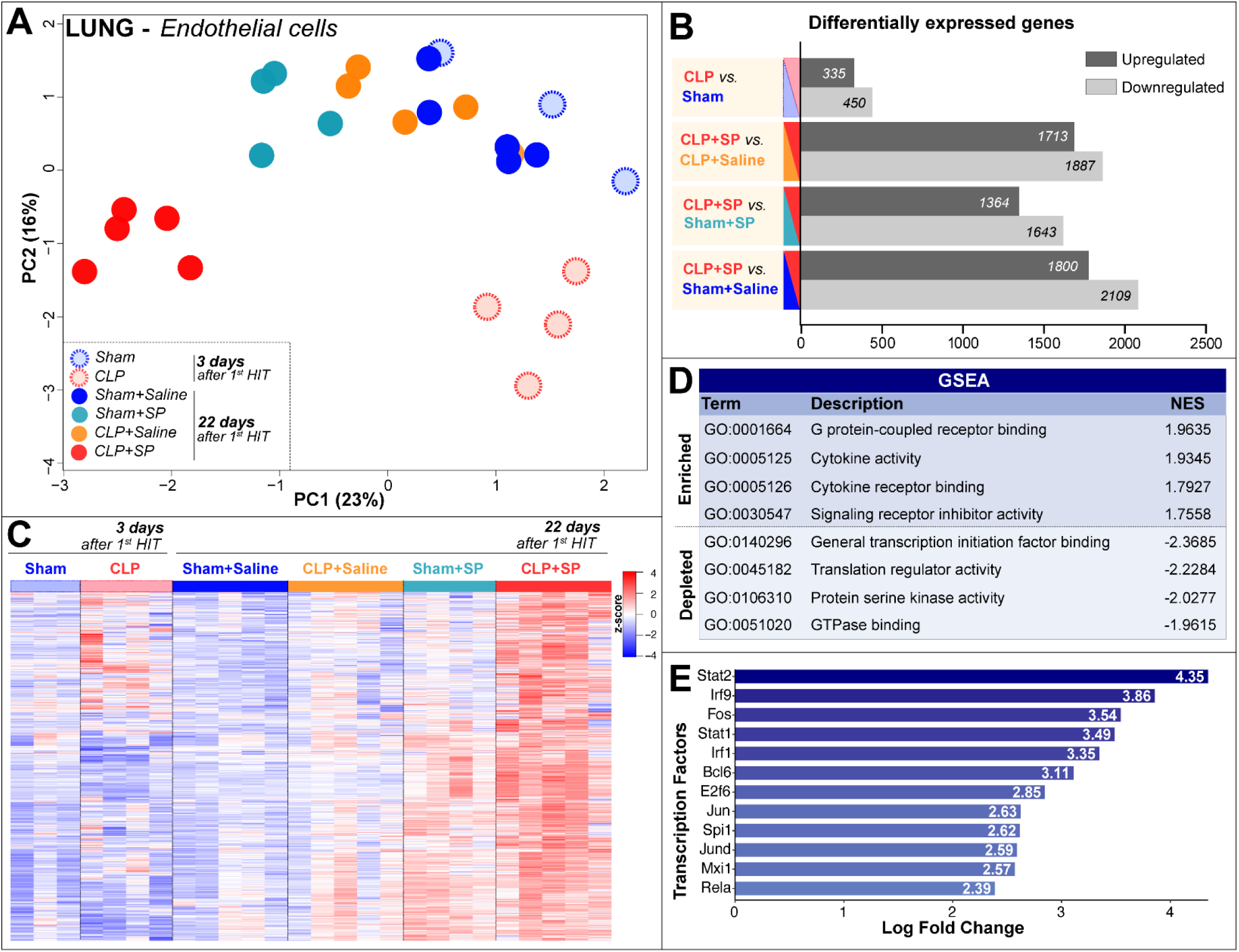
Lung Endothelium Exhibits Amplified Transcriptional Activation After Secondary Infection. **(A)** Principal component analysis (PCA) of lung endothelial RNA-seq data shows clear separation of CLP+SP samples from all other groups, including CLP-only (day 3) and all day 22 controls, indicating distinct transcriptional reprogramming upon secondary infection. **(B)** Barplot of differentially expressed genes (DEGs) highlights substantial upregulation and downregulation specifically in CLP+SP mice across comparisons. **(C)** Heatmap of amplified genes demonstrates robust transcriptional activation in CLP+SP mice **(D)** Gene set enrichment analysis (GSEA) reveals strong enrichment of cytokine activity, receptor binding, and G protein– coupled receptor signaling terms, alongside suppression of general transcription factor and kinase-related pathways (WebGestalt). **(E)** Transcription factor activity analysis identifies high predicted activation of Stat, Irf and AP-1 complex members, supporting a coordinated transcriptional network driving inflammatory memory in the lung endothelium (DoRothEA, NES > 2).

Differential gene expression analysis confirmed this robust transcriptional response, with CLP+SP endothelial cells showing the highest number of significantly upregulated and downregulated genes compared to all other experimental groups (**Fig. 3B, supplemental table 1**).

To specifically identify genes uniquely and robustly amplified in CLP+SP lungs, we used a selection criteria requiring genes to be significantly upregulated (adjusted p-value < 0.05 and log₂FC fold change > 0.5) in CLP+SP versus all other comparisons (Sham+SP, Sham+saline). The resulting gene set, visualized by heatmap, showed 1385 genes uniquely upregulated in the CLP+SP group. These included prominent interferon-stimulated genes (e.g., Irf7, Ifit1, Ifit2, Ifit3, Ifi204, Ifi207, Rsad2, Oasl1, Oasl2, Mx1), inflammatory cytokines and chemokines (e.g., Ccl2, Ccl5, Cxcl9, Cxcl10, Cxcl2, Il1b, Tnf), pathogen recognition receptors (Tlr2, Tlr6), complement components (C1ra, C1rb, C3, Cfb, C4a), and critical regulators of cell survival and inflammatory signaling (Icam1, Vcam1, Cd14, Stat2, Irf9, Nfkbia, Nfkbiz, Tnfaip3) (**Fig. 3C**). These genes reflect a robust endothelial activation, with potential implications for enhanced leukocyte recruitment, sustained inflammation, and endothelial dysfunction.

Gene Set Enrichment Analysis (GSEA) from the amplified genes on the CLP+SP further supported these findings, highlighting significant enrichment in molecular functions associated with cytokine activity, cytokine receptor binding, signaling receptor activity, and response to microbial ligands such as lipopolysaccharide (**Fig. 3D**). Such functional enrichment reinforces the mechanistic link between endothelial transcriptional reprogramming and amplified inflammatory responses upon secondary infectious challenges. Additionally, transcription factor activity analysis on the amplified genes revealed significant activation of key inflammatory transcriptional regulators in CLP+SP endothelial cells, prominently involving Stat2, Irf9, Stat1, Fos, Irf1, Bcl6, and NF-κB family members (Rela, Rel, Nfkb1) (**Fig. 3E**). Underscoring a coordinated inflammatory and interferon-driven response, central to establishing endothelial inflammatory memory and heightened susceptibility to secondary insults in post-sepsis lungs. Immunofluorescence further confirms endothelial dysfunction and immune cell infiltration, as evidenced by increased P-selectin (**Fig. 4**).

**Fig 4.**
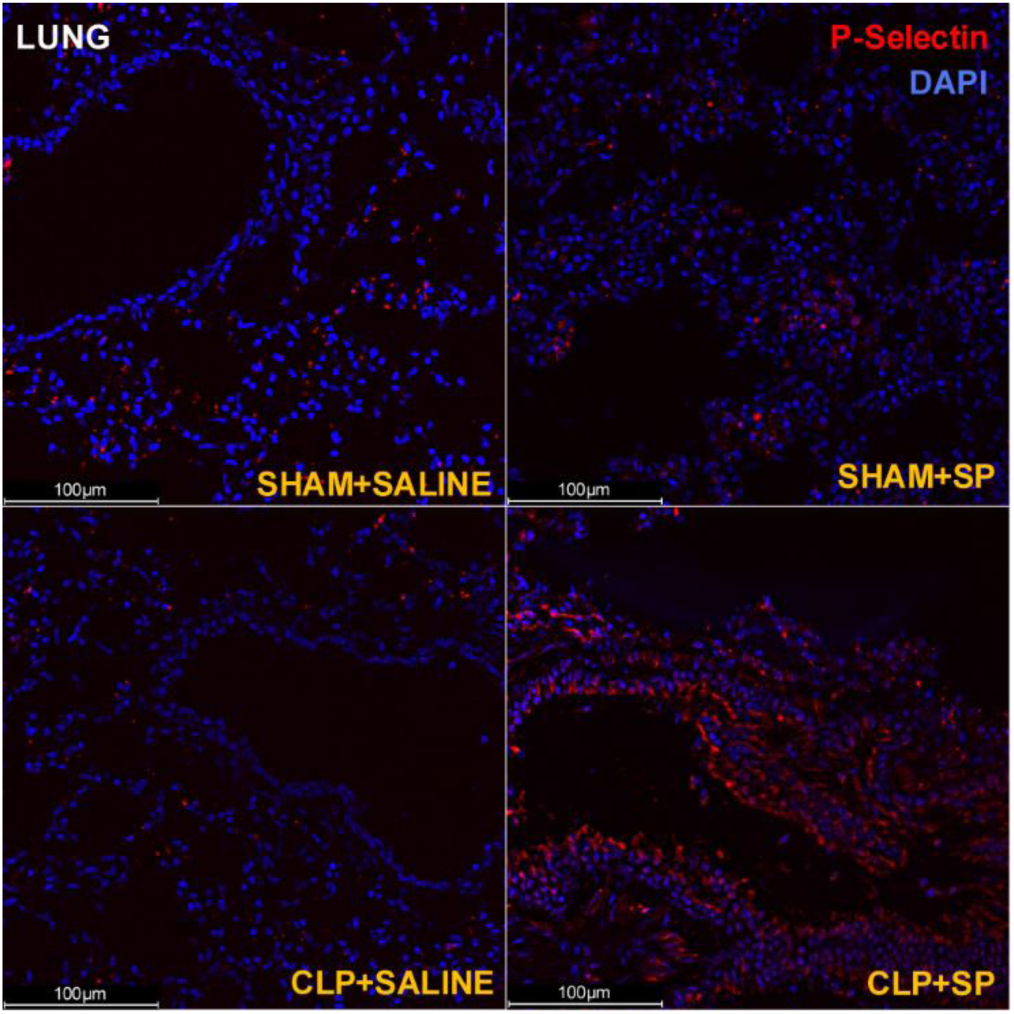
Immunofluorescence analysis of P-selectin expression in lung tissue following secondary challenge. Representative lung sections from SHAM+SALINE, SHAM+SP, CLP+SALINE, and CLP+SP mice were stained for P-selectin (red) and nuclei (DAPI, blue) to assess endothelial activation at day 22 post-first hit. P-selectin expression was minimal in SHAM groups, with modest induction in CLP+SALINE. In contrast, CLP+SP lungs showed robust P-selectin staining localized to vascular structures, indicating heightened endothelial activation following the two-hit injury. Scale bars, 100 μm.

### Transcriptional Reprogramming of Kidney Endothelium in Post-Sepsis Mice

Given that the kidney is a clinically relevant site of long-term dysfunction in sepsis survivors (Flannery, Li et al. 2021), we next hypothesized whether the transcriptional priming observed in lung endothelium also extends to the renal vasculature. We profiled ECs isolated from kidneys harvested at the same experimental conditions as the lungs. PCA distinctly separated kidney ECs from the CLP+SP group from all other conditions, including acute CLP (day 3), CLP+saline, sham+SP, and sham+saline controls (**Fig. 5A**). Differential expression analysis further confirmed this transcriptional reprogramming, identifying substantial numbers of significantly differentially expressed genes uniquely elevated in CLP+SP endothelial cells compared to CLP alone, highlighting an amplified endothelial response following secondary infection (**Fig. 5B, supplemental table 2**).

**Fig 5.**
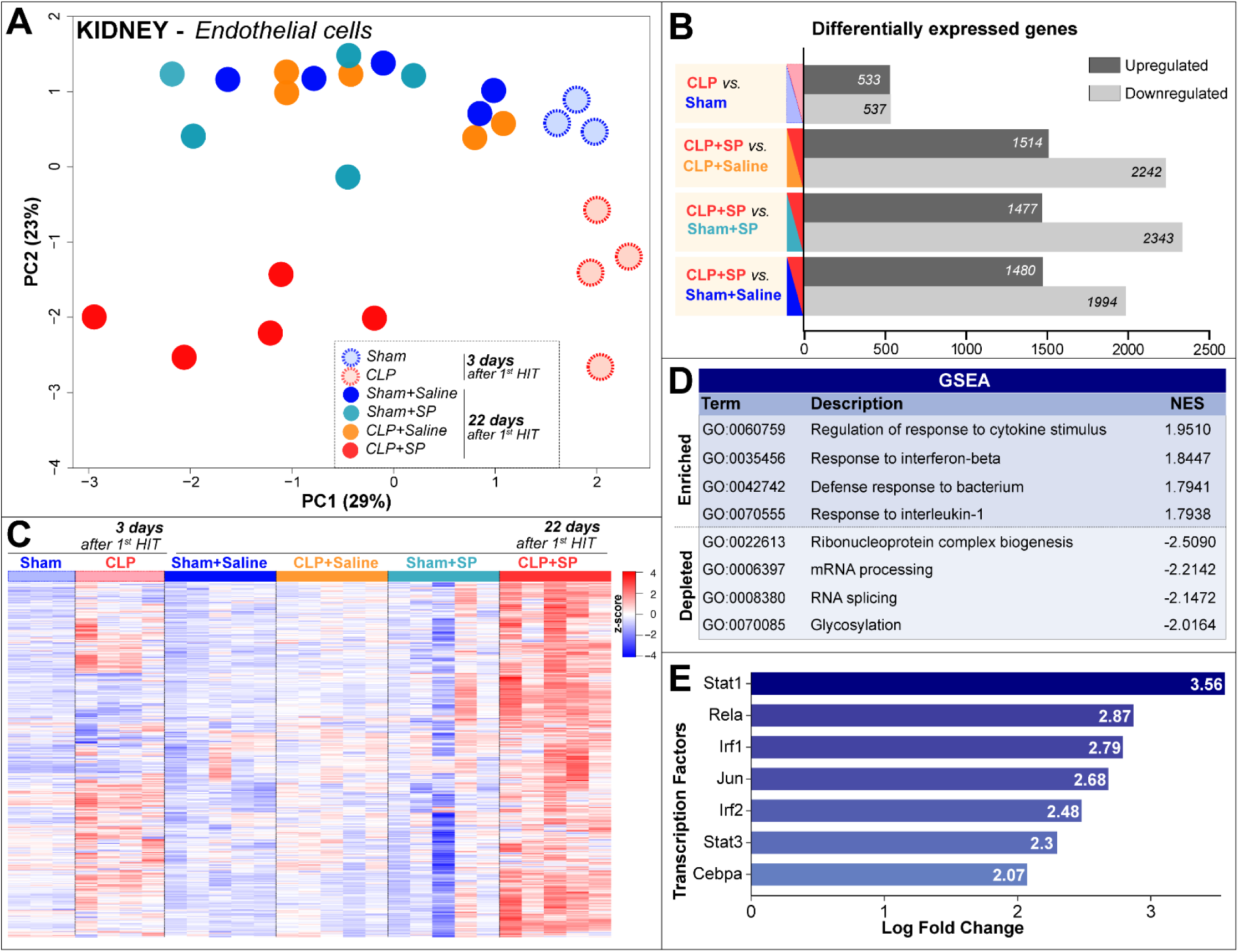
Kidney Endothelium Demonstrates Robust and Distinct Transcriptional Priming Following Secondary Challenge. **(A)** PCA of kidney endothelial RNA-seq data reveals clear segregation of CLP+SP samples from all other experimental groups, including CLP alone (day 3) and all controls at day 22, indicating unique transcriptional reprogramming following secondary challenge. **(B)** DEG highlights extensive transcriptional changes in CLP+SP. **(C)** Heatmap of 1,385 amplified genes in CLP+SP samples highlights a distinct and coordinated activation of inflammatory, immune-regulatory, and metabolic genes relative to other conditions. **(D)** GSEA identifies significant enrichment of cytokine signaling, interferon responses, and bacterium defense pathways in CLP+SP kidneys (WebGestalt) **(E)** Transcription factor activity analysis reveals elevated activation of Stat1, Rela (NF-κB), Irf and Jun, suggesting a coordinated transcriptional network underlying endothelial inflammatory memory in the kidney (DoRothEA, NES > 2).

Using same criteria as the lung, we identified a set of 748 genes amplified in the kidney endothelium of CLP+SP mice, visualized by heatmap with separation from other groups (**Fig. 5C**). This amplified gene set included critical inflammatory mediators and endothelial activation markers (Icam1, Vcam1, Sele), interferon-stimulated genes (Ifit3, Irgm1, Oasl2, Ifitm3), antigen presentation molecules (H2-K1, H2-Q4, Tap1), cytokine and chemokine signaling components (Csf2rb, Il18bp, Socs1, Socs3), complement cascade components (C1ra, C1rb, C3, Cfb), and key regulatory factors of inflammatory and immune signaling (Tnfaip3, Nfkbiz, Stat3, Irf1, Irf9, Junb). These findings underscore a transcriptional amplification consistent with persistent endothelial inflammation and activation following secondary infection. GSEA reinforced these observations, demonstrating significant enrichment of pathways involved in cytokine responses, interferon-beta signaling, leukocyte-mediated immunity, and antigen processing, consistent with heightened endothelial immunological activation (**Fig. 5D**).

Transcription factor activity analysis indicated significant activation of key inflammatory and immune-related transcription factors, notably Stat1, NF-κB (Rela), Irf1, Jun, Irf2, Stat3, and C/EBP family members (Cebpa, Cebpb, Cebpd) (**Fig. 5E**). These transcription factors likely orchestrate the amplified and persistent transcriptional responses observed in the kidney endothelium of sepsis survivors exposed to a secondary insult. Consistent with the transcriptional enrichment of endothelial adhesion molecules observed in the kidney RNA-seq data, immunofluorescence staining revealed a marked increase in P-selectin expression in renal vasculature of CLP+SP mice, supporting sustained endothelial activation and inflammatory priming following secondary challenge (**Fig. 6**).

**Figure 6.**
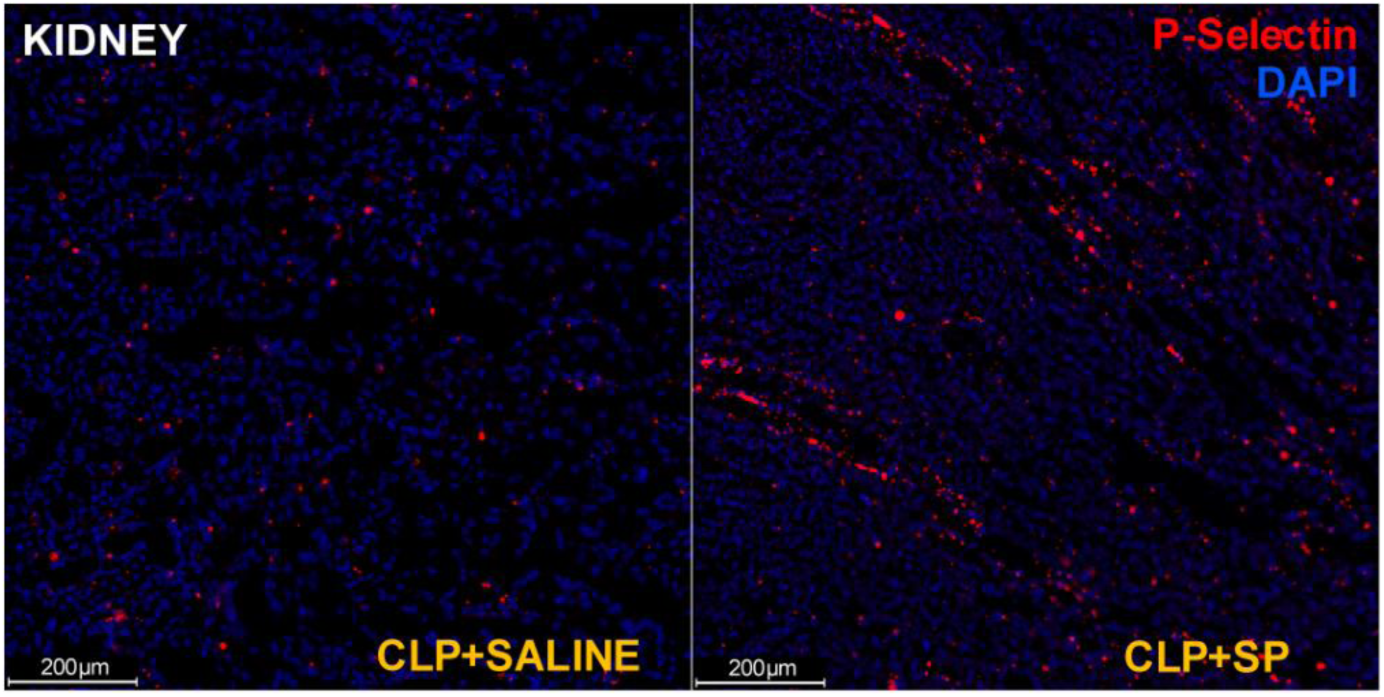
P-selectin immunofluorescence reveals enhanced endothelial activation in kidneys of CLP+SP mice. Kidney sections from CLP+SALINE and CLP+SP groups were stained for P-selectin (red) and nuclei (DAPI, blue) at day 22 post-sepsis. CLP+SP kidneys exhibited markedly increased P-selectin expression along vascular structures compared to CLP+SALINE, consistent with persistent endothelial activation and inflammatory priming following secondary Streptococcus pneumoniae challenge. Scale bars, 200 μm.

These data demonstrate that prior sepsis establishes enduring endothelial transcriptional priming in the kidney, leading to exaggerated responses upon secondary infectious challenges, potentially underlying the enhanced susceptibility to chronic organ dysfunction frequently observed in sepsis survivors.

### Shared and Organ-specific Endothelial Inflammatory Memory in Post-Sepsis Dysfunction

Our study highlights a molecular framework underlying endothelial reprogramming following repeated inflammatory insults, a set of 301 genes was significantly amplified in endothelial cells from both lung and kidney, underscoring common endothelial responses to secondary inflammatory challenges. Notably, this shared signature includes genes critical for complement activation (C1ra, C1rb, C3, Cfb), innate immune regulation (Irgm1, Ifitm3), endothelial activation and leukocyte adhesion (Icam1), and negative inflammatory feedback (Tnfaip3, Serping1). This conserved transcriptional response positions endothelial cells as active mediators of sustained inflammation and vascular injury across multiple organs, highlighting a common mechanism underlying post-sepsis vulnerability (**Supplemental Fig 2A-B, supplemental table 3**).

Lung ECs exhibited a robust interferon-driven phenotype, characterized by upregulation of antiviral response genes (Ifit1, Ifit2, Rsad2, Ifi204), transcriptional regulators (Irf7, Irf8), and leukocyte recruitment chemokines (Cxcl9, Cxcl10, Ccl2, Ccl7). Concurrent increases in PRRs (Tlr2, Zbp1, Clec5a) suggest heightened surveillance for microbial products, aligning with the lung’s role as a frontline barrier against airborne pathogens. In contrast, kidney ECs displayed transcriptional enrichment in genes associated with metabolic adaptation and immune modulation. Genes such as Nampt, Slc3a2, Slc46a3, and Gars1 point to elevated nutrient transport and energy metabolism, potentially supporting endothelial endurance and recovery within the renal microenvironment. Additional upregulation of barrier-regulatory and immune-regulatory genes (Ecscr, Emp1, H2-K1, Lgals9) underscores the kidney’s unique immunovascular niche **(Supplemental Fig 2A)**

To mechanistically resolve the transcriptional control of these programs, we performed transcription factor activity inference using DoRothEA. Shared responses were dominated by canonical inflammatory regulators such as STAT1, NFKB1, and RELA, while organ-specific signatures revealed TF divergence. Lung memory responses preferentially activated JUN/FOS, SPI1, and STAT1, while kidney-specific responses were shaped by TEAD1 (**Supplemental Fig 2B**). These data provide evidence that endothelial inflammatory memory is both transcriptionally imprinted and contextually shaped by tissue environment, offering opportunities for organ-specific therapeutic targeting in post-sepsis complications.

### Persistent and Worsened Kidney Dysfunction in Post-Sepsis Mice

While our transcriptomic analyses revealed robust and organ-specific endothelial activation, molecular changes alone do not fully capture the physiological consequences of this inflammatory memory. To determine whether these transcriptional reprogramming events translate into measurable functional outcomes, we next assessed glomerular filtration rate (GFR), an indicator of renal performance. We hypothesized that sepsis survivors would exhibit long-lasting renal vulnerability that could be exacerbated by a second inflammatory hit.

Despite apparent clinical recovery by day 13, CLP survivors exhibited significantly reduced GFR compared to sham-operated controls, indicating subclinical kidney dysfunction (**Fig. 7A**). This impairment persisted beyond the recovery phase and was further exacerbated following the secondary *SP* challenge. By day 21, one day after the second hit, GFR in the CLP+SP group dropped significantly below all other groups, including CLP+saline and SHAM+SP (**Fig. 7B**). These findings reveal that renal vulnerability persists in sepsis survivors and is unmasked by a secondary infectious insult, paralleling clinical observations where post-septic patients show heightened risk for kidney injury and long-term renal complications.

**Fig 7.**
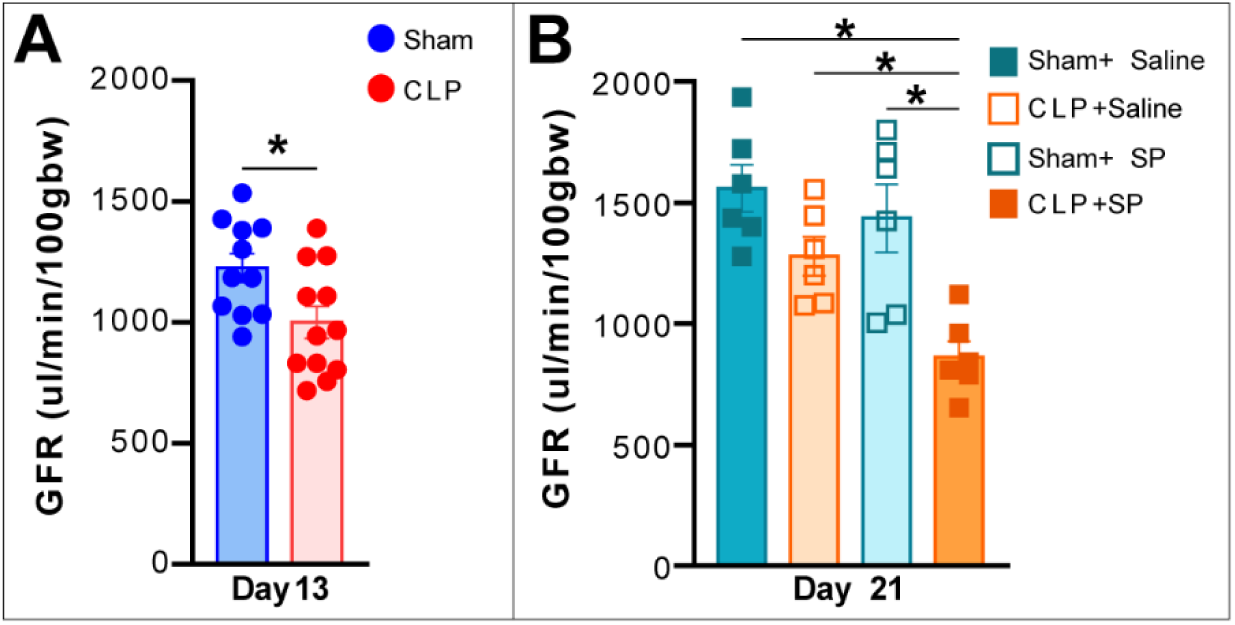
Persistent and Worsened Kidney Dysfunction Following Secondary Challenge. **(A)** Glomerular filtration rate (GFR) measured by FITC-sinistrin clearance at day 13 post-CLP, demonstrating persistent subclinical kidney impairment in CLP mice **(B)** day 21 (one day post-secondary SP challenge), indicating exacerbated dysfunction uniquely in CLP+SP mice. Data represent mean±SEM; *p<0.05 by one-way ANOVA with Tukey’s multiple comparisons.

### Endothelial Cells Display Amplified ICAM-1 Activation Following Secondary Challenge

While endothelial transcriptomic profiling revealed robust activation signatures in sepsis survivors, we sought to determine whether these molecular changes were reflected in cell-surface phenotypes relevant to vascular inflammation. Given the central role of adhesion molecules in leukocyte recruitment, we next assessed endothelial expression of ICAM-1 and P-selectin in the kidney following secondary challenge. Although the overall frequency of endothelial cells (CD45⁻/CD31^+^) remained unchanged across groups (**Fig. 8A**), CLP mice challenged with *SP* exhibited an increase in intensity of ICAM-1 expression on kidney endothelial cells (**Fig. 8B**). P-selectin expression, in contrast, remained stable across conditions (**Fig. 8C**). These findings point to a primed endothelial phenotype in sepsis survivors, characterized by heightened ICAM-1 responsiveness (Sessler, Windsor et al. 1995, Kaur, Hussain et al. 2021, He, Duan et al. 2024).

**Fig 8.**
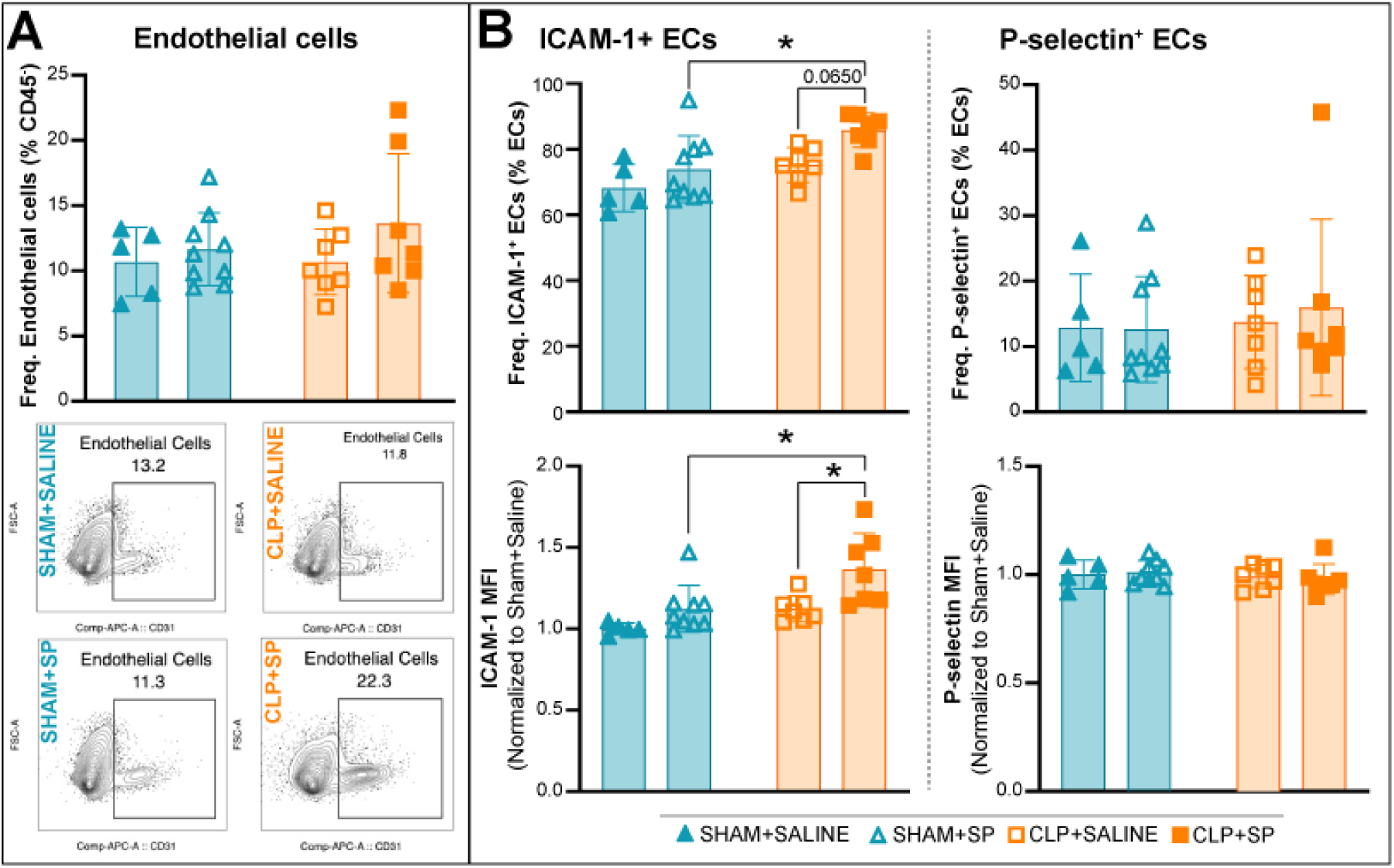
Kidney Endothelium Exhibits Enhanced ICAM-1 Expression Following Secondary Challenge in Sepsis Survivors. **(A)** Flow cytometry quantification of endothelial cells (CD45⁻CD31⁺) in kidney tissue shows no significant difference in overall endothelial cell frequency across experimental groups on day 22. **(B)** CLP+SP mice display a significant increase in both the frequency and surface expression (mean fluorescence intensity, MFI) of ICAM-1⁺ endothelial cells compared to all other groups. **(C)** No significant changes were observed in the frequency or MFI of P-selectin⁺ endothelial cells. Data are presented as mean ± SEM. Statistical comparisons were performed using one-way ANOVA with Tukey’s multiple comparisons.; *p < 0.05.

### Heightened Myeloid Activation and Immature Neutrophil Expansion in Post-Sepsis Kidneys

To investigate cellular immune responses underlying kidney dysfunction in post-sepsis mice, we profiled kidney immune cell composition following secondary *SP* challenge. Total kidney cellularity remained comparable across groups, with CD45⁺ hematopoietic cells consistently representing a minority fraction (**Fig. 9A**). However, notable shifts in myeloid cell populations emerged specifically in sepsis survivors re-challenged with *SP*. The frequency of neutrophils significantly increased in the kidneys of CLP+SP mice compared to sham controls (**Fig. 9B**). There was a pronounced elevation in Ly6C^hi inflammatory monocytes in CLP+SP mice, reflecting enhanced recruitment and infiltration of these pro-inflammatory cells into the kidney tissue following secondary insult (**Fig. 9B**).

**Fig 9.**
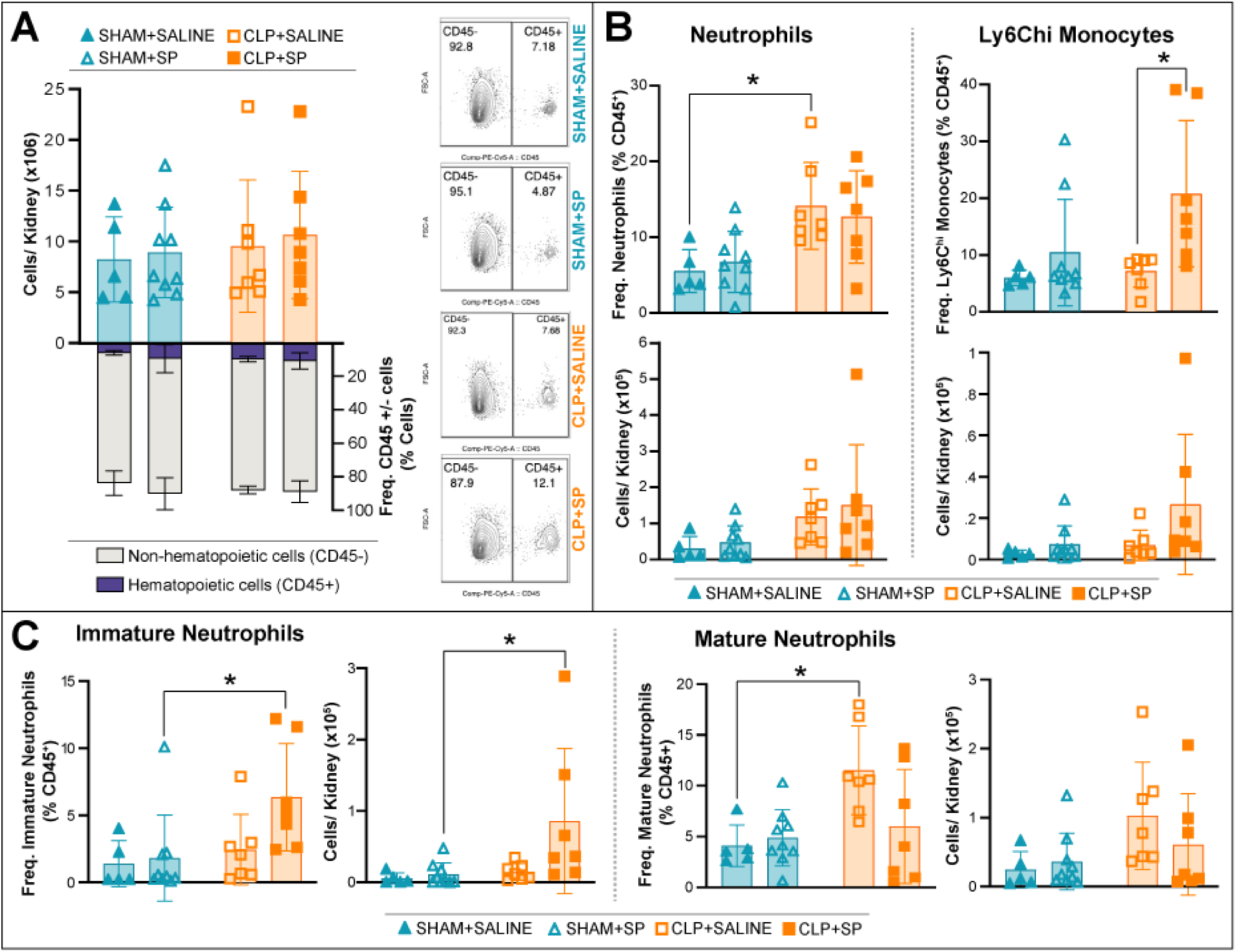
Expansion of Inflammatory Monocytes and Immature Neutrophils in Post-Sepsis Kidneys Following Secondary Challenge. **(A)** Total kidney cellularity (top) and the proportion of CD45⁺ hematopoietic cells (bottom) remained unchanged across experimental groups on day 22. Representative contour plots show CD45⁺ gating. **(B)** CLP+SP kidneys exhibited a significant increase in the frequency and absolute number of total inflammatory Ly6C^hi monocytes (CD11b⁺Ly6C^hi) compared to all other groups. **(C)** The myeloid compartment in CLP+SP kidneys showed a selective expansion of immature neutrophils (CD101⁻), both in frequency and number, whereas mature neutrophils (CD101⁺) were not significantly altered. These findings indicate a shift toward emergency myelopoiesis and persistent immune activation following sepsis and secondary infection. Data are presented as mean ± SEM; statistical significance determined by one-way ANOVA with Tukey’s post hoc test; *p < 0.05.

Further characterization of neutrophil subsets revealed a marked accumulation of immature neutrophils (CD11b⁺Ly6G⁺CD101⁻) in the CLP+SP group, representing a substantial proportion of total neutrophils (**Fig. 9C**). Conversely, mature neutrophil frequencies (CD101⁺) remained stable, indicating a selective expansion of immature neutrophils following secondary infection (**Fig. 9C**). Representative contour plots further illustrate this disproportionate increase in immature neutrophils and inflammatory monocytes specifically in the CLP+SP kidneys (**Supplemental Figure 3**). Underscoring a maladaptive immune response characterized by heightened recruitment of inflammatory monocytes and expansion of immature neutrophils in post-sepsis kidneys. This altered myeloid landscape likely contributes to persistent inflammation, impaired resolution, and ultimately worsened kidney dysfunction following secondary infectious insults.

### HUVEC – Investigating Endothelial Inflammatory Memory Through Transcriptional Profiling

To understand the long-term impact of inflammatory priming on endothelial cells, we used an integrated transcriptomic (RNA-seq) and epigenomic (ATAC-seq) approach in human umbilical vein endothelial cells (HUVECs) treated sequentially with IL-6 and lipopolysaccharide (LPS). Cells were initially stimulated with IL-6 plus its soluble receptor (IL-6+R, 200 ng/mL) for 72 hours, followed by a 48-hour washout to simulate resolution. Subsequently, cells were challenged with a mild dose of LPS (1 µg/mL) for 6 hours, modeling a secondary infection (**Fig. 10A**). Through integrative RNA-seq and ATAC-seq analyses, we aim to elucidate how initial inflammatory priming may stably reprogram the endothelium, laying the foundation for exacerbated secondary responses.

**Fig 10.**
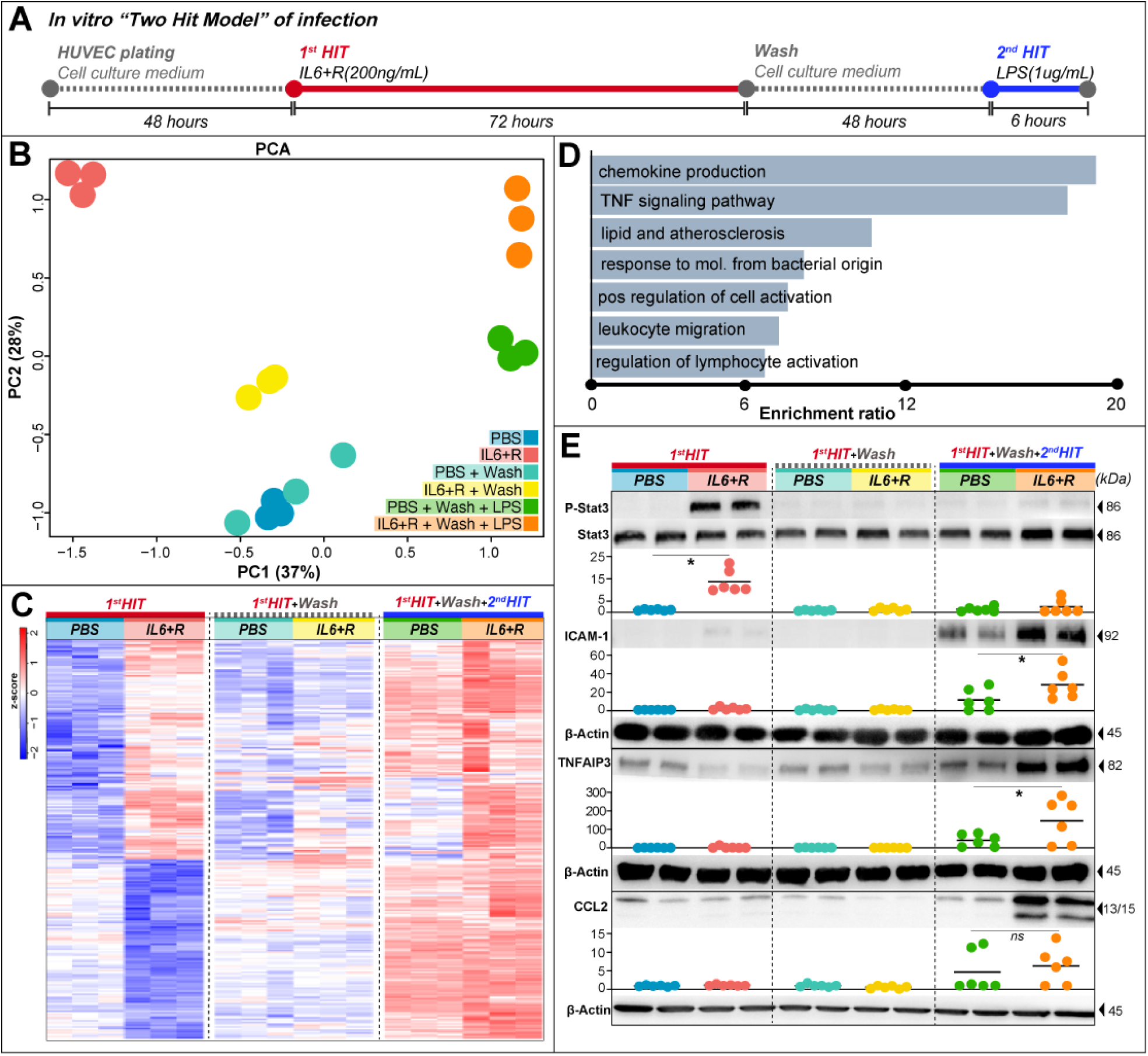
Transcriptional reprogramming in endothelial inflammatory memory model. **(A)** Schematic representation of the in vitro endothelial two-hit inflammatory model, HUVECs treated with IL-6+R (200 ng/mL) for 72h (1st hit), followed by 48h washout and subsequent LPS stimulation (1 µg/mL, 6h, 2nd hit). **(B)** PCA reveals distinct transcriptomic profiles between treatment groups, highlighting separation of cells receiving IL-6 priming followed by LPS (IL-6+R+Wash+LPS). **(C)** Heatmap depicting significant gene expression changes across treatment groups, illustrating robust transcriptional amplification specifically after the second inflammatory stimulus. **(D)** GO analysis (WebGestalt) identifies significantly enriched inflammatory pathways activated after sequential inflammatory challenges. **(E)** Western blot quantification demonstrates sustained STAT3 phosphorylation during the first hit and increased ICAM-1 and TNFAIP3 protein expression following the second hit. β-actin served as loading control; Data presented as mean ± SEM; statistical significance determined by one-way ANOVA with Tukey’s post hoc test; *p < 0.05.

PCA from transcriptomic data distinctly clustered IL-6-treated samples (red) separately from PBS controls (blue), with IL-6-primed cells challenged with LPS (orange) forming another unique cluster, clearly demonstrating persistent transcriptional alterations due to initial IL-6 exposure (**Fig. 10B**). Following the secondary LPS challenge, we identified 86 genes exhibiting significantly amplified expression compared to the primary IL-6 stimulus alone (**Fig. 10C, Supplemental Table 4**). Genes were considered amplified if they met a log_2_FC threshold > 0.5 and adjusted p-value < 0.05 in both the initial IL-6 exposure and the subsequent LPS challenge. The wash condition genes were included with no significant differential expression observed between IL-6+Wash and PBS+Wash groups (adj. p-value > 0.05), indicating transcriptional recovery between challenges. Amplification was defined by a higher log_2_FC in the second challenge (IL-6+Wash+LPS) compared to the primary IL-6 exposure or LPS alone in the second hit, indicating an enhanced transcriptional response upon re-stimulation.

Gene ontology enrichment analyses highlighted key inflammatory and immune regulatory pathways including chemokine production, TNF signaling, and leukocyte migration (**Fig. 10D**). Notably, genes within the chemokine production pathway (GO:0032602), such as IL6, TLR2, LGALS9, HMOX1, APOD, IL1RL1, and TICAM1, emerged as central mediators of inflammation. Among these, IL6 and TLR2 were prominent, recurring across multiple enriched pathways, thus emphasizing their critical roles in orchestrating broad-spectrum inflammatory responses and endothelial activation. Additionally, enriched genes involved in TNF signaling (e.g., CCL2, CXCL2, ICAM1, NFKBIA, TNFAIP3) highlighted the interplay between chemokine production and NF-κB signaling modulation. Particularly, TNFAIP3, known for its role as a negative regulator of inflammation (Catrysse, Vereecke et al. 2014), was upregulated following IL-6 priming and exhibited further induction upon subsequent LPS challenge, indicative of active feedback regulation.

Validation at the protein level reinforced these transcriptional insights. Western blot analyses demonstrated activation of the IL-6–STAT3 signaling pathway, evidenced by increased phosphorylated STAT3 (P-STAT3) after initial IL-6 stimulation, which returned to baseline after the washout period. Importantly, ICAM-1 protein was minimally expressed following the initial IL-6 treatment but significantly elevated after the subsequent LPS challenge, specifically in IL-6-primed cells. The persistent elevation of TNFAIP3 protein further substantiated active feedback regulation aimed at controlling inflammatory magnitude (**Fig. 10E**).

To test whether the order of inflammatory stimuli influences the induction of endothelial memory, we performed an inverted two-hit experiment in which LPS was used as the primary stimulus followed by IL-6+R re-challenge. Unlike the canonical IL-6-first paradigm, this inverted model did not lead to amplified transcriptional responses upon IL-6 exposure. Expression of key inflammatory genes, including IL6, IL8, CCL2, ICAM1, COX2, and TNFAIP3, remained comparable to single-stimulus controls, suggesting that TLR4-driven priming via LPS is insufficient to establish a memory state responsive to IL-6 reactivation. These findings indicate that IL-6–STAT3 signaling may be specifically required during the initial stimulus to program chromatin and transcriptional responsiveness characteristic of inflammatory memory (**Supplemental Figure 4**). We demonstrated that initial IL-6 exposure imprints endothelial cells with a durable transcriptional signature, fundamentally altering their responsiveness and promoting an amplified inflammatory phenotype upon secondary stimulus. This enhanced understanding of endothelial inflammatory memory provides critical insights into the sustained vascular dysfunction observed in chronic inflammatory conditions such as sepsis.

### ATAC-seq Analysis Reveals Stable Chromatin Remodeling and Correlation with Transcriptional Changes

To determine if IL-6–induced inflammatory memory involves persistent epigenetic remodeling, we performed ATAC-seq in HUVECs exposed to IL-6+R or PBS for 72 h, followed by a 48-hour wash period. Genome-wide analysis revealed widespread chromatin accessibility changes in response to IL-6. An elbow plot ranking peaks by their log₂ fold change normalized to standard error (log₂FC/SEM) showed a clear bifurcation, with strongly upregulated (open) peaks at one extreme and downregulated (closed) peaks at the other, highlighting the most dynamic regions of the epigenome (**Fig. 11A**). In total, 56,010 peaks became more accessible (open), while 45,428 peaks exhibited decreased accessibility (closed) compared to PBS controls after Il6 treatment for 72 hours (**Fig. 11B**). Importantly, nearly half of the IL-6–induced open regions (26,747 peaks) remained accessible even after the cytokine was removed and cells were washed for 48 hours, indicating persistent chromatin accessibility (**Fig. 11B**). This subset of stably open peaks likely represents epigenetically imprinted regulatory elements, consistent with a transcriptionally poised state underlying endothelial inflammatory memory, a list of all 26,747 peaks with chromosome, genomic start and end and a list of all position overlapping with genes and number of peaks for each gene are show at **Supplemental table 5**.

**Fig 11.**
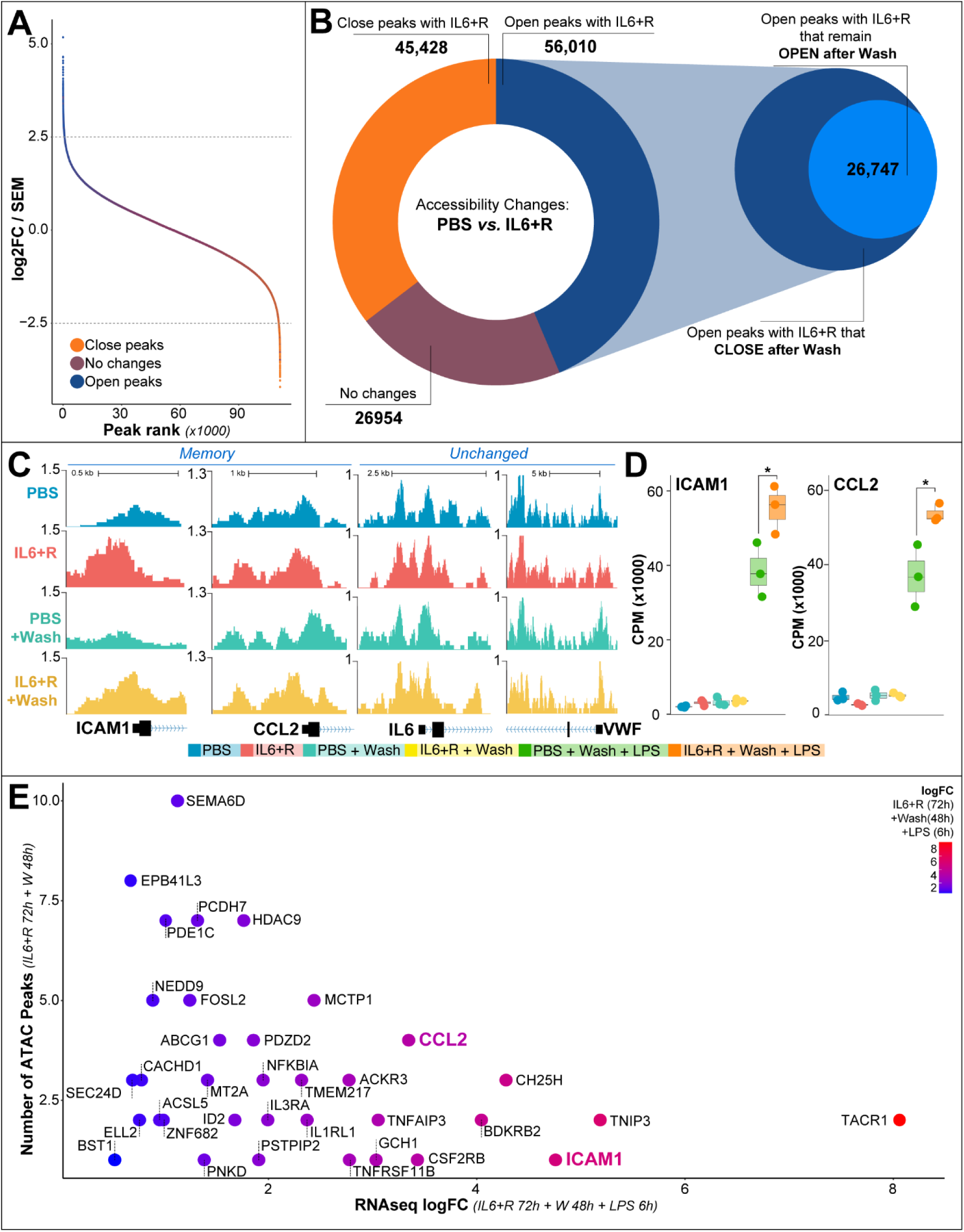
IL-6-induced persistent chromatin remodeling correlates with transcriptional memory in endothelial cells. **(A)** Elbow plot ranking peaks by log₂ fold-change normalized to standard error (log₂FC/SEM), indicating the most dynamically altered accessible chromatin regions after IL-6+R exposure. **(B)** Donut plot illustrates total chromatin accessibility changes: 56,010 open peaks and 45,428 closed peaks induced by IL-6+R; 27,171 peaks remained persistently open even after 48-hour washout. **(C-D)** Genome browser tracks and RNA-seq quantification highlight persistent accessibility and elevated expression of inflammatory memory-associated genes (ICAM1, CCL2), contrasting with unchanged genes (IL6, VWF). **(E)** Scatterplot identifies key genes with concordant increases in ATAC-seq peak number and transcriptional amplification after the second inflammatory stimulus. Color intensity indicates magnitude of transcriptional change.

We further characterized these persistent chromatin changes at the gene level, integrating chromatin accessibility with RNA-seq data. A set of genes, including ICAM1 and CCL2, displayed persistent open chromatin peaks post-wash, correlating directly with elevated transcriptional responses upon a secondary inflammatory stimulus (**Fig. 10C–D**). Genes, such as IL6 and VWF, exhibited no substantial changes in chromatin accessibility after IL-6 treatment or subsequent wash, indicating selective rather than global epigenetic remodeling.

Expanding beyond ICAM1 and CCL2, we identified 56 genes exhibiting robust increases in both chromatin accessibility and transcription, such as FOSL2, MAP3K8, NFKBIA, TNFAIP3, SEMA6D, EPB41L3, PDE1C, PCDH7, and HDAC9(**Fig. 11E**). Several of these genes had multiple sustained open chromatin peaks, indicative of strong regulatory reinforcement. Notably, genes such as FOSL2 and NFKBIA are well-established targets regulated by the AP-1/JunB transcription factor complex, critical in mediating inflammatory signaling and chromatin remodeling (Fujioka, Niu et al. 2004, Vierbuchen, Ling et al. 2017, Renoux, Stellato et al. 2020, Yukawa, Jagannathan et al. 2020). AP-1 transcription factors, particularly JunB, heterodimerize with FOS family members (including FOSL2) to regulate enhancer activation and subsequent inflammatory gene transcription (Renoux, Stellato et al. 2020, Karakaslar, Katiyar et al. 2023). MAP3K8 (TPL-2), TNFAIP3 (A20), and NFKBIA (IκBα) are pivotal regulators within inflammatory pathways, tightly controlled by NF-κB and AP-1 interactions, providing a critical feedback mechanism that modulates endothelial inflammation and immune response intensity (Yu, Lin et al. 2020, Nanou, Bourbouli et al. 2021). Collectively, our results reveal that IL-6 exposure induces a stable and functionally relevant chromatin landscape, characterized by persistent regulatory accessibility at inflammatory response genes. This epigenetic imprinting provides a mechanistic foundation for heightened transcriptional responsiveness upon subsequent inflammatory encounters, potentially contributing to the prolonged endothelial dysfunction and increased susceptibility to secondary infections observed in post-septic patients.

### JunB Is Required for IL-6–Mediated Chromatin and Transcriptional Remodeling

To understand the mechanism driving IL-6–induced inflammatory memory, we investigated the role of JunB, a core component of the AP-1 transcription factor complex previously implicated in enhancer activation and inflammatory gene regulation. Given our integrative chromatin and RNA-seq data identifying multiple JunB-regulated targets—including ICAM1, CCL2, FOSL2, and TNFAIP3—as persistently open and transcriptionally amplified following IL-6 exposure, we hypothesized that JunB is a critical driver of this epigenetic reprogramming.

We performed siRNA-mediated knockdown of JunB (siJunB) in HUVECs treated with IL-6+R for 72 hours to mimic the first-hit inflammatory exposure. We confirmed efficient JunB silencing at the transcript levels (**Fig. 12A**). JunB knockdown led to transcriptional suppression of IL-6–inducible genes.

**Fig 12.**
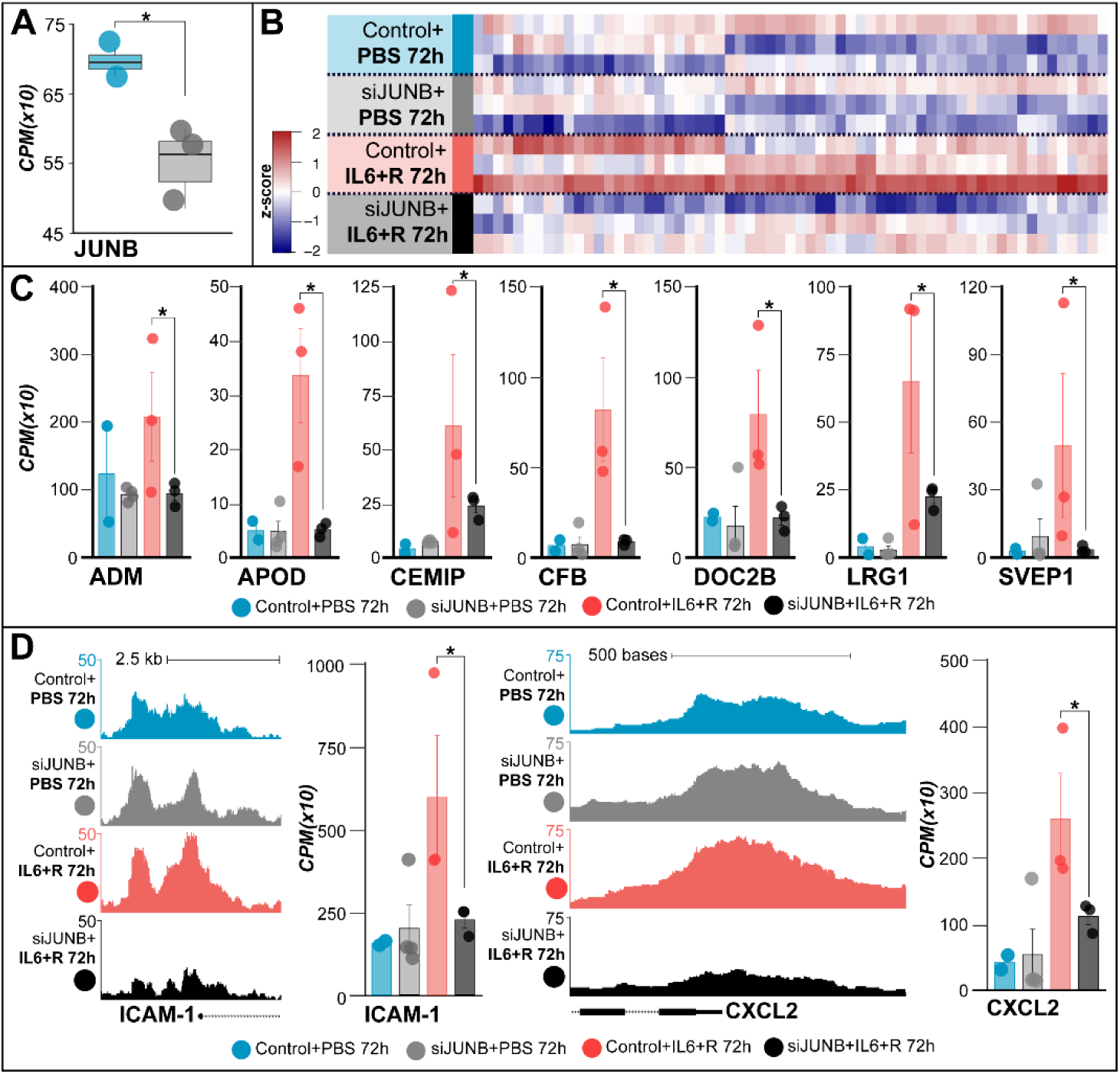
JunB is critical for IL-6-mediated transcriptional memory in endothelial cells. **(A)** RNA-seq quantification showing effective siRNA-mediated JunB knockdown in HUVECs. **(B)** Heatmap comparing gene expression profiles reveals significant suppression of IL-6–induced inflammatory gene amplification upon JunB knockdown. **(C)** RNA-seq validation identifies selected inflammatory memory-associated genes (ADM, APOD, CEMIP, CFB, DOC2B, LRG1, SVEP1) whose amplified by the two hit model is significantly reduced by siJunB after IL6+R for 72 hours. **(D)** ATAC-seq tracks and corresponding RNA expression of ICAM-1 and CXCL2 demonstrate JunB-dependent chromatin accessibility and transcriptional upregulation after IL-6+R stimulation.

Analysis revealed a downregulation expression across 63 genes, showing a JunB-sensitive gene set, with multiple transcripts displaying attenuated induction following IL-6 treatment in the absence of JunB (**Fig. 12B**). Specifically, genes such as ADM, APOD, CEMIP, CFB, DOC2B, LRG1, and SVEp1, all amplified by the second hit (**Fig. 10C**), were significantly reduced upon JunB silencing (**Fig. 12C**). Over representation analysis showed the genes are associated with TNF signaling pathway (CEBPB, ICAM1, E-Selectin), IL signaling pathway (CEBPB, CSF3, IKBKE), vascular homeostasis (HLA, ICAM1, MPZ, E-selectin), supporting JunB’s role in shaping a proinflammatory endothelial phenotype. Chromatin accessibility and gene expression levels of canonical inflammatory targets ICAM1 and CXCL2 were also markedly diminished in siJunB-treated cells, both at the epigenetic and transcript level (**Fig. 12D**), consistent with JunB-dependent enhancer activation and transcriptional priming.

Having established that JunB is critical for orchestrating chromatin remodeling and transcriptional activation during the initial IL-6 exposure, we next sought to determine whether JunB remains necessary for the amplified transcriptional response during a subsequent inflammatory challenge. To mimic the “second hit,” we silenced JunB using siRNA and stimulated endothelial cells with LPS for 6 hours, modeling acute exposure in a primed context. Western blot and qPCR confirmed effective JunB knockdown across both PBS and LPS-stimulated groups **(Fig. 13A)**. JunB deficiency significantly reduced the expression of canonical inflammatory response genes such as ICAM1 and COX2 upon LPS stimulation **(Fig. 13B)**, indicating that JunB remains functionally required for the transcriptional amplification observed during secondary challenge. Interestingly, expression of IL6, IL8, TNFAIP3, and FOS was not significantly altered by JunB silencing, suggesting that only a subset of LPS-responsive genes are JunB-dependent at this stage.

**Fig 13.**
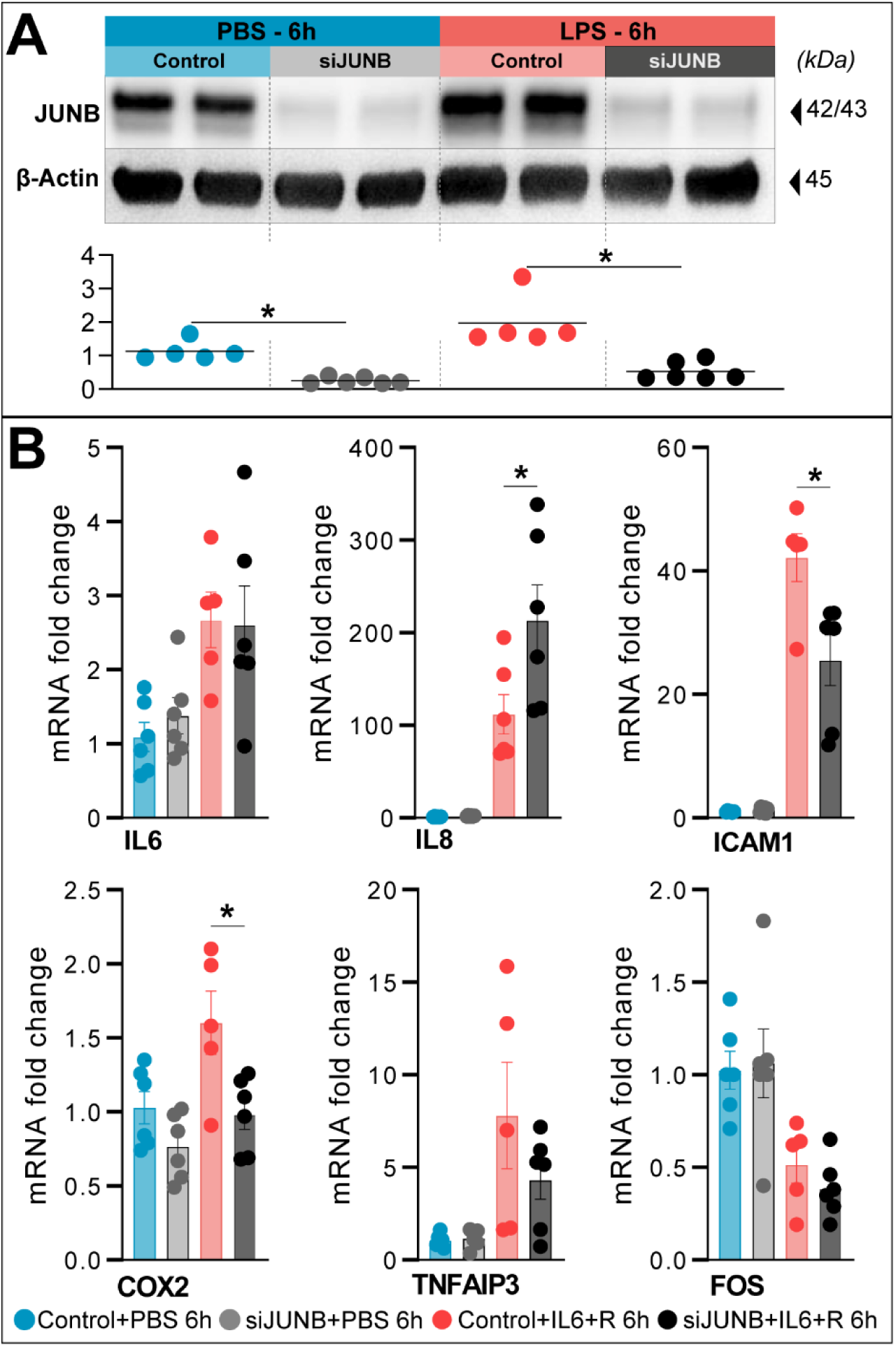
JunB mediates amplified endothelial responses to LPS challenge. **(A)** Western blot and quantification demonstrate effective JunB knockdown in endothelial cells exposed to PBS or LPS (6h). β-actin used as loading control. **(B)** qRT-PCR analysis shows significant reduction in LPS-induced expression of inflammatory genes ICAM1, COX2, and IL8 in JunB-knockdown cells, with limited effects on IL6, TNFAIP3, and FOS. Data presented as mean ± SEM; *p<0.05, ANOVA with Tukey’s post-hoc test.

These results reveal a sustained regulatory role for JunB beyond initial stimulus exposure. JunB is not only essential for priming endothelial chromatin and transcriptional landscapes but also continues to mediate selective gene activation during secondary inflammatory responses. This dual-phase requirement positions JunB as a key transcriptional node in the endothelial inflammatory memory circuit, offering a mechanistic explanation for persistent endothelial dysfunction in the post-sepsis setting.

## Discussion

Our study reveals a previously unrecognized mechanism of endothelial inflammatory memory wherein transient inflammatory exposure induces durable chromatin remodeling at multiple critical inflammatory and regulatory loci—including not only canonical pro-inflammatory genes like ICAM1 and CCL2 but also key transcriptional regulators such as FOSL2, MAP3K8, NFKBIA, and TNFAIP3. This sustained epigenetic reprogramming, driven by IL-6–STAT3 signaling and reinforced by AP-1/JunB transcription factor complexes, primes endothelial cells for exaggerated transcriptional responses upon subsequent inflammatory stimuli, significantly amplifying systemic cytokine release, leukocyte recruitment, and ultimately resulting in profound susceptibility to secondary infections.

These findings are highly relevant to secondary infection risk after acute inflammation episodes, as sepsis with higher risk of mortality, with up to 30% of patients succumbing to secondary infections(Jarczak, Kluge et al. 2021). Our findings provide mechanistic insight into how ECs retain a transcriptionally poised state following inflammatory exposure, driven by persistent chromatin accessibility at key inflammatory loci. ATAC-seq analyses revealed that IL-6 stimulation leads to widespread chromatin remodeling, with a subset of regions remaining accessible even after cytokine withdrawal—so-called memory peaks. These persistent open chromatin regions were associated with genes involved in cytokine signaling, leukocyte recruitment, and immune regulation, implicating them as central nodes in the endothelial memory program. Importantly, elevated expression of ICAM1 and CCL2 has been independently linked to endothelial dysfunction in sepsis and worse clinical outcomes (Shapiro, Schuetz et al. 2010, Jia, Xu et al. 2022, Zhang, Wang et al. 2023, He, Duan et al. 2024), while downregulation of TNFAIP3 is associated with sustained inflammatory signaling (Catrysse, Vereecke et al. 2014). These epigenetic features may therefore contribute to the chronic vascular activation and increased vulnerability to secondary infections observed in sepsis survivors.

The endothelial inflammatory memory concept parallels observations in innate immune cells, where trained immunity is characterized by long-lasting epigenetic changes that enhance transcriptional responsiveness to secondary insults (Drummer, Saaoud et al. 2021, Flores-Gomez, Bekkering et al. 2021, Riksen, Bekkering et al. 2023). In monocytes, for example, β-glucan stimulation induces H3K27ac and H3K4me1 enrichment at inflammatory gene enhancers, establishing a poised chromatin state conducive to rapid gene activation (Saeed, Quintin et al. 2014).

Work by Sohrabi et al. (2022) has further advanced this concept by demonstrating that activation of the Liver X Receptor (LXR), a central regulator of lipid metabolism, induces a trained immunity phenotype in human monocytes through coordinated epigenetic and metabolic reprogramming. In their study, LXR agonists not only enhanced the proinflammatory effects of classical training stimuli such as BCG but were also capable of independently inducing long-term inflammatory activation. This was accompanied by increased histone acetylation at inflammatory gene promoters, elevated acetyl-CoA levels, and dependence on the IL-1β signaling, mechanisms that mirror known features of trained immunity in macrophages. Interestingly, inhibition of fatty acid synthesis paradoxically intensified this proinflammatory memory, suggesting a finely tuned metabolic balance in memory formation. These findings highlight how non-infectious stimuli, through metabolic rewiring, can stably remodel the epigenome to produce durable inflammatory phenotypes (Sohrabi, Sonntag et al. 2020), a principle we now extend to endothelial cells in the context of sepsis-induced vascular dysfunction Consistent with this, a study revealed that epithelial stem cells also retain a chromatin-based memory of prior inflammatory exposures, characterized by persistent open chromatin at memory domains, enrichment of activating histone marks (H3K4me1, H3K27ac), and sustained occupancy of transcription factors like JUN. In their model, STAT3 cooperated with AP-1 components (FOS and JUN) to establish these memory domains during the primary inflammatory response, after which JUN remained bound, acting as a scaffold to recruit FOS rapidly upon secondary stimulation (Larsen, Cowley et al. 2021). This coordinated handoff enables a broad and rapid recall response, even to stimuli distinct from the initial insult. Our data support a parallel mechanism in endothelial cells, where IL-6-induced STAT3 activation primes chromatin accessibility, and JunB functions as an amplifier upon subsequent LPS challenge, bridging stimulus-specific initiation with general stress-responsive transcriptional amplification.

Specific in ECs, previous study showed transient high-glucose exposure induces prolonged proinflammatory signaling through Set7-mediated methylation (Okabe, Orlowski et al. 2012) Similarly, oxidized phospholipids associated with Lp(a) have been shown to reprogram ECs toward a glycolytic, proinflammatory state (Schnitzler, Hoogeveen et al. 2020), further supporting the concept that ECs can undergo stable transcriptional and metabolic reprogramming in response to stimuli. This aligns with observations in vascular smooth muscle cells (vSMCs), where reduced levels of the repressive histone mark H3K9me2 facilitate enhanced binding of NFκB and AP-1 transcription factors at inflammation-responsive promoters such as IL6 and MMP3, thereby promoting sustained proinflammatory gene expression in response to inflammatory stimuli (Harman, Dobnikar et al. 2019). Together, these studies reinforce a universal model of inflammatory memory across both immune and non-immune cells, where early stimulus-specific transcription factors initiate chromatin remodeling, and enables rapid and amplified responses upon re-challenge.

Our findings suggest that IL-6–induced inflammatory memory may contribute to the persistent endothelial activation seen in sepsis survivors and help explain why these patients remain vulnerable to secondary infections despite apparent clinical recovery (Dolmatova, Wang et al. 2021). Interestingly, when we reversed the order of inflammatory stimuli—initially priming endothelial cells with LPS followed by IL-6+R, this amplification of the transcriptional response was not observed. A critical methodological detail is that our IL-6 priming conditions included the soluble IL-6 receptor (sIL-6R), forming an IL-6/sIL-6R complex known to robustly activate STAT3 signaling via trans-signaling mechanisms. The absence of sIL-6R during initial LPS priming suggests that the IL-6/sIL-6R trans-signaling pathway might be specifically required to establish a durable chromatin accessibility state and subsequent transcriptional amplification. This distinction underscores the critical role of IL-6 trans-signaling in generating endothelial memory and provides clinical insight into the use of targeted therapeutic interventions.

Early blockade of IL-6 signaling using agents such as tocilizumab, currently in use for cytokine storm syndromes (Barrett 2024), may limit the establishment of epigenetic memory if applied during the acute phase of inflammation. Specifically, early intervention with Tocilizumab, a monoclonal antibody targeting IL-6 signaling showed the patients receiving early intervention post-sepsis induction, exhibited the most favorable molecular and highest survival rate when compared with the group receiving the latest treatment post-sepsis induction(Tavaci, Halici et al. 2025). One hypothesis can be by the initial epigenetic priming step if administered during the acute phase of inflammation. Consequently, biomarkers such as ICAM1 and CCL2 not only reflect the presence of endothelial memory but also hold translational potential as predictors of adverse long-term outcomes, aiding in patient stratification and therapeutic decision-making(Kaur, Hussain et al. 2021, He, Duan et al. 2024, Peng, Wang et al. 2024). Moreover, the fact that these molecular changes are recapitulated in primary human ECs highlights their clinical relevance and translational potential. Therapeutically, these insights open new avenues for intervention.

In addition, our data implicate JunB, a component of the activator protein-1 (AP-1) transcription factor complex, as a key regulator that links persistent chromatin accessibility to amplified transcriptional responses during secondary inflammatory challenges. AP-1 is a heterodimeric complex composed primarily of members of the JUN (e.g., JUN, JunB, JunD) and FOS (e.g., c-FOS, FOSL1, FOSL2) families, and plays a central role in coordinating transcriptional programs in response to stress, cytokines, and pathogen-associated molecular patterns(Fujioka, Niu et al. 2004, Vierbuchen, Ling et al. 2017, Karakaslar, Katiyar et al. 2023, Song, Lian et al. 2023). Upon inflammatory stimulation, such as IL-6 or LPS exposure, AP-1 binds to enhancer elements marked by histone modifications like H3K27ac, promoting the expression of genes involved in cytokine production, adhesion molecule expression, and immune cell recruitment(Borghini, Hibberd et al. 2018, Herrera-Uribe, Liu et al. 2020). Specifically, JunB has been shown to act not only as a transcriptional activator but also as a pioneer-like factor that modulates chromatin accessibility in concert with other stress-responsive transcription factors(Ito, Yamauchi et al. 2001, Ndlovu, Van Lint et al. 2009, Papavassiliou and Musti 2020).

Given its central role in orchestrating inflammation-related gene networks, targeting AP-1 activity—either through direct inhibition of JunB or upstream signaling pathways (e.g., MAPK, JAK/STAT)—represents a promising strategy to disrupt the pathological transcriptional amplification associated with endothelial memory. Moreover, epigenetic co-factors that facilitate AP-1-mediated enhancer activation, such as p300/CBP or BRD4, are druggable, making HDAC and BET inhibitors attractive candidates to reverse maladaptive epigenetic states in post-sepsis vasculature. Our findings thus establish JunB/AP-1 as a mechanistic and potentially therapeutic node linking transient inflammatory stimuli to long-term endothelial reprogramming.

While our study significantly advances the understanding of endothelial inflammatory memory, several limitations should be acknowledged as avenues for future exploration. First, the current in vivo model was limited to a follow-up of approximately three weeks post-sepsis, precluding insights into the very long-term persistence of endothelial epigenetic remodeling and the chronic impact on organ dysfunction. Future studies extending survival and functional assessment beyond this period could better delineate the durability and clinical implications of endothelial memory. Additionally, our primary human endothelial model relied on HUVECs, which represent macrovascular endothelium and may not fully capture the intrinsic functional and transcriptional heterogeneity of microvascular endothelial cells, critical mediators of inflammation-induced organ dysfunction in sepsis. Subsequent studies utilizing tissue-specific microvascular endothelial cell models or single-cell epigenomic profiling would further strengthen the translational relevance of these findings. Finally, although our results robustly implicate JunB/AP-1 as central mediators of chromatin remodeling and transcriptional amplification, definitive validation via targeted genetic models, such as endothelial-specific JunB knockout mice, would provide deeper mechanistic insight and confirm causality. Such approaches could illuminate the therapeutic potential and safety profiles of targeting JunB or related chromatin regulators to mitigate persistent vascular inflammation post-sepsis.

## Notes

### Competing Interest Statement

The authors have declared no competing interest.

### Funding

- American Heart Association: 24CDA1268714

## References

1. Augustin, H. G. and G. Y. Koh (2024). “A systems view of the vascular endothelium in health and disease.” Cell 187(18): 4833–4858.

2. Badia, I. M. P., J. Velez Santiago, J. Braunger, C. Geiss, D. Dimitrov, S. Muller-Dott, P. Taus, A. Dugourd, C. H. Holland, R. O. Ramirez Flores and J. Saez-Rodriguez (2022). “decoupleR: ensemble of computational methods to infer biological activities from omics data.” Bioinform Adv 2(1): vbac016.

3. Biswas, N., A. Bahr, J. Howard, J. L. Bonin, R. Grazda and K. C. MacNamara (2024). “Survivors of polymicrobial sepsis are refractory to G-CSF-induced emergency myelopoiesis and hematopoietic stem and progenitor cell mobilization.” Stem Cell Reports 19(5): 639–653.

4. Borghini, L., M. Hibberd and S. Davila (2018). “Changes in H3K27ac following lipopolysaccharide stimulation of nasopharyngeal epithelial cells.” BMC Genomics 19(1): 969.

5. Bossardi Ramos, R., N. Martino, D. Chuy, S. Lu, M. X. G. Zuo, U. Balasubramanian, I. Di John Portela, P. A. Vincent and A. P. Adam (2023). “Shock drives a STAT3 and JunB-mediated coordinated transcriptional and DNA methylation response in the endothelium.” J Cell Sci.

6. Catrysse, L., L. Vereecke, R. Beyaert and G. van Loo (2014). “A20 in inflammation and autoimmunity.” Trends Immunol 35(1): 22–31.

7. Chen, H. and P. C. Boutros (2011). “VennDiagram: a package for the generation of highly-customizable Venn and Euler diagrams in R.” BMC Bioinformatics 12: 35.

8. Chen, Y., L. Chen, A. T. L. Lun, P. L. Baldoni and G. K. Smyth (2025). “edgeR v4: powerful differential analysis of sequencing data with expanded functionality and improved support for small counts and larger datasets.” Nucleic Acids Res 53(2).

9. Dolmatova, E. V., K. Wang, R. Mandavilli and K. K. Griendling (2021). “The effects of sepsis on endothelium and clinical implications.” Cardiovasc Res 117(1): 60–73.

10. Drummer, C. t., F. Saaoud, Y. Shao, Y. Sun, K. Xu, Y. Lu, D. Ni, D. Atar, X. Jiang, H. Wang and X. Yang (2021). “Trained Immunity and Reactivity of Macrophages and Endothelial Cells.” Arterioscler Thromb Vasc Biol 41(3): 1032–1046.

11. Elizarraras, J. M., Y. Liao, Z. Shi, Q. Zhu, A. R. Pico and B. Zhang (2024). “WebGestalt 2024: faster gene set analysis and new support for metabolomics and multi-omics.” Nucleic Acids Res 52(W1): W415–W421.

12. Fernandez, S., M. Palomo, P. Molina, M. Diaz-Ricart, G. Escolar, A. Tellez, F. Segui, H. Ventosa, S. Torramade-Moix, M. Rovira, E. Carreras, J. M. Nicolas and P. Castro (2021). “Progressive endothelial cell damage in correlation with sepsis severity. Defibrotide as a contender.” J Thromb Haemost 19(8): 1948–1958.

13. Flannery, A. H., X. Li, N. L. Delozier, R. D. Toto, O. W. Moe, J. Yee and J. A. Neyra (2021). “Sepsis-Associated Acute Kidney Disease and Long-term Kidney Outcomes.” Kidney Med 3(4): 507–514 e501.

14. Flores-Gomez, D., S. Bekkering, M. G. Netea and N. P. Riksen (2021). “Trained Immunity in Atherosclerotic Cardiovascular Disease.” Arterioscler Thromb Vasc Biol 41(1): 62–69.

15. Fujioka, S., J. Niu, C. Schmidt, G. M. Sclabas, B. Peng, T. Uwagawa, Z. Li, D. B. Evans, J. L. Abbruzzese and P. J. Chiao (2004). “NF-kappaB and AP-1 connection: mechanism of NF-kappaB-dependent regulation of AP-1 activity.” Mol Cell Biol 24(17): 7806–7819.

16. Garcia-Alonso, L., C. H. Holland, M. M. Ibrahim, D. Turei and J. Saez-Rodriguez (2019). “Benchmark and integration of resources for the estimation of human transcription factor activities.” Genome Res 29(8): 1363–1375.

17. Grandi, F. C., H. Modi, L. Kampman and M. R. Corces (2022). “Chromatin accessibility profiling by ATAC-seq.” Nat Protoc 17(6): 1518–1552.

18. Harman, J. L., L. Dobnikar, J. Chappell, B. G. Stokell, A. Dalby, K. Foote, A. Finigan, P. Freire-Pritchett, A. L. Taylor, M. D. Worssam, R. R. Madsen, E. Loche, A. Uryga, M. R. Bennett and H. F. Jorgensen (2019). “Epigenetic Regulation of Vascular Smooth Muscle Cells by Histone H3 Lysine 9 Dimethylation Attenuates Target Gene-Induction by Inflammatory Signaling.” Arterioscler Thromb Vasc Biol 39(11): 2289–2302.

19. He, J., M. Duan and H. Zhuang (2024). “ICAM1 and VCAM1 are associated with outcome in patients with sepsis: A systematic review and meta-analysis.” Heliyon 10(21): e40003.

20. Herrera-Uribe, J., H. Liu, K. A. Byrne, Z. F. Bond, C. L. Loving and C. K. Tuggle (2020). “Changes in H3K27ac at Gene Regulatory Regions in Porcine Alveolar Macrophages Following LPS or PolyIC Exposure.” Front Genet 11: 817.

21. Hotchkiss, R. S., L. L. Moldawer, S. M. Opal, K. Reinhart, I. R. Turnbull and J. L. Vincent (2016). “Sepsis and septic shock.” Nat Rev Dis Primers 2: 16045.

22. Ince, C., P. R. Mayeux, T. Nguyen, H. Gomez, J. A. Kellum, G. A. Ospina-Tascon, G. Hernandez, P. Murray, D. De Backer and A. X. Workgroup (2016). “The Endothelium in Sepsis.” Shock 45(3): 259–270.

23. Inghammar, M., A. Linder, M. Lengquist, A. Frigyesi, H. Wetterberg, J. Sunden-Cullberg and A. Nilsson (2024). “Long-term Mortality and Hospital Readmissions Among Survivors of Sepsis in Sweden: A Population-Based Cohort Study.” Open Forum Infect Dis 11(7): ofae331.

24. Ito, T., M. Yamauchi, M. Nishina, N. Yamamichi, T. Mizutani, M. Ui, M. Murakami and H. Iba (2001). “Identification of SWI.SNF complex subunit BAF60a as a determinant of the transactivation potential of Fos/Jun dimers.” J Biol Chem 276(4): 2852–2857.

25. Jarczak, D., S. Kluge and A. Nierhaus (2021). “Sepsis-Pathophysiology and Therapeutic Concepts.” Front Med (Lausanne) 8: 628302.

26. Jia, P., S. Xu, X. Wang, X. Wu, T. Ren, Z. Zou, Q. Zeng, B. Shen and X. Ding (2022). “Chemokine CCL2 from proximal tubular epithelial cells contributes to sepsis-induced acute kidney injury.” Am J Physiol Renal Physiol 323(2): F107–F119.

27. Joffre, J., J. Hellman, C. Ince and H. Ait-Oufella (2020). “Endothelial Responses in Sepsis.” Am J Respir Crit Care Med 202(3): 361–370.

28. Kaneki, M. (2017). “Metabolic Inflammatory Complex in Sepsis: Septic Cachexia as a Novel Potential Therapeutic Target.” Shock 48(6): 600–609.

29. Karakaslar, E. O., N. Katiyar, M. Hasham, A. Youn, S. Sharma, C. H. Chung, R. Marches, R. Korstanje, J. Banchereau and D. Ucar (2023). “Transcriptional activation of Jun and Fos members of the AP-1 complex is a conserved signature of immune aging that contributes to inflammaging.” Aging Cell 22(4): e13792.

30. Kaur, S., S. Hussain, K. Kolhe, G. Kumar, D. M. Tripathi, A. Tomar, P. Kale, A. Narayanan, C. Bihari, M. Bajpai, R. Maiwall, E. Gupta and S. K. Sarin (2021). “Elevated plasma ICAM1 levels predict 28-day mortality in cirrhotic patients with COVID-19 or bacterial sepsis.” JHEP Rep 3(4): 100303.

31. Landt, S. G., G. K. Marinov, A. Kundaje, P. Kheradpour, F. Pauli, S. Batzoglou, B. E. Bernstein, P. Bickel, J. B. Brown, P. Cayting, Y. Chen, G. DeSalvo, C. Epstein, K. I. Fisher-Aylor, G. Euskirchen, M. Gerstein, J. Gertz, A. J. Hartemink, M. M. Hoffman, V. R. Iyer, Y. L. Jung, S. Karmakar, M. Kellis, P. V. Kharchenko, Q. Li, T. Liu, X. S. Liu, L. Ma, A. Milosavljevic, R. M. Myers, P. J. Park, M. J. Pazin, M. D. Perry, D. Raha, T. E. Reddy, J. Rozowsky, N. Shoresh, A. Sidow, M. Slattery, J. A. Stamatoyannopoulos, M. Y. Tolstorukov, K. P. White, S. Xi, P. J. Farnham, J. D. Lieb, B. J. Wold and M. Snyder (2012). “ChIP-seq guidelines and practices of the ENCODE and modENCODE consortia.” Genome Res 22(9): 1813–1831.

32. Larsen, S. B., C. J. Cowley, S. M. Sajjath, D. Barrows, Y. Yang, T. S. Carroll and E. Fuchs (2021). “Establishment, maintenance, and recall of inflammatory memory.” Cell Stem Cell 28(10): 1758–1774 e1758.

33. Law, C. W., Y. Chen, W. Shi and G. K. Smyth (2014). “voom: Precision weights unlock linear model analysis tools for RNA-seq read counts.” Genome Biol 15(2): R29.

34. Lawrence, M., W. Huber, H. Pages, P. Aboyoun, M. Carlson, R. Gentleman, M. T. Morgan and V. J. Carey (2013). “Software for computing and annotating genomic ranges.” PLoS Comput Biol 9(8): e1003118.

35. Li, H., B. Handsaker, A. Wysoker, T. Fennell, J. Ruan, N. Homer, G. Marth, G. Abecasis, R. Durbin and S. Genome Project Data Processing (2009). “The Sequence Alignment/Map format and SAMtools.” Bioinformatics 25(16): 2078–2079.

36. Liao, Y., G. K. Smyth and W. Shi (2019). “The R package Rsubread is easier, faster, cheaper and better for alignment and quantification of RNA sequencing reads.” Nucleic Acids Res 47(8): e47.

37. Lu, S., I. D. John Portela, N. Martino, R. B. Ramos, A. E. Salinero, R. M. Smith, K. L. Zuloaga and A. P. Adam (2024). “A transient brain endothelial translatome response to endotoxin is associated with mild cognitive changes post-shock in young mice.” bioRxiv.

38. Lu, Y., Y. Sun, K. Xu, Y. Shao, F. Saaoud, N. W. Snyder, L. Yang, J. Yu, S. Wu, W. Hu, J. Sun, H. Wang and X. Yang (2022). “Editorial: Endothelial cells as innate immune cells.” Front Immunol 13: 1035497.

39. Martino, N., R. Bossardi Ramos, D. Chuy, L. Tomaszek and A. P. Adam (2022). “SOCS3 limits TNF and endotoxin-induced endothelial dysfunction by blocking a required autocrine interleukin-6 signal in human endothelial cells.” Am J Physiol Cell Physiol 323(2): C556–C569.

40. Martino, N., R. B. Ramos, S. Lu, K. Leyden, L. Tomaszek, S. Sadhu, G. Fredman, A. Jaitovich, P. A. Vincent and A. P. Adam (2021). “Endothelial SOCS3 maintains homeostasis and promotes survival in endotoxemic mice.” JCI Insight 6(14).

41. Naik, S. and E. Fuchs (2022). “Inflammatory memory and tissue adaptation in sickness and in health.” Nature 607(7918): 249–255.

42. Nanou, A., M. Bourbouli, S. Vetrano, U. Schaeper, S. Ley and G. Kollias (2021). “Endothelial Tpl2 regulates vascular barrier function via JNK-mediated degradation of claudin-5 promoting neuroinflammation or tumor metastasis.” Cell Rep 35(8): 109168.

43. Nascimento, D. C., P. R. Viacava, R. G. Ferreira, M. A. Damaceno, A. R. Pineros, P. H. Melo, P. B. Donate, J. E. Toller-Kawahisa, D. Zoppi, F. P. Veras, R. S. Peres, L. Menezes-Silva, D. Caetite, A. E. R. Oliveira, I. M. S. Castro, G. Kauffenstein, H. I. Nakaya, M. C. Borges, D. S. Zamboni, D. M. Fonseca, J. A. R. Paschoal, T. M. Cunha, V. Quesniaux, J. Linden, F. Q. Cunha, B. Ryffel and J. C. Alves-Filho (2021). “Sepsis expands a CD39(+) plasmablast population that promotes immunosuppression via adenosine-mediated inhibition of macrophage antimicrobial activity.” Immunity 54(9): 2024–2041 e2028.

44. Ndlovu, M. N., C. Van Lint, K. Van Wesemael, P. Callebert, D. Chalbos, G. Haegeman and W. Vanden Berghe (2009). “Hyperactivated NF-kappaB and AP-1 transcription factors promote highly accessible chromatin and constitutive transcription across the interleukin-6 gene promoter in metastatic breast cancer cells.” Mol Cell Biol 29(20): 5488–5504.

45. Netea, M. G., J. Dominguez-Andres, L. B. Barreiro, T. Chavakis, M. Divangahi, E. Fuchs, L. A. B. Joosten, J. W. M. van der Meer, M. M. Mhlanga, W. J. M. Mulder, N. P. Riksen, A. Schlitzer, J. L. Schultze, C. Stabell Benn, J. C. Sun, R. J. Xavier and E. Latz (2020). “Defining trained immunity and its role in health and disease.” Nat Rev Immunol 20(6): 375–388.

46. Niec, R. E., A. Y. Rudensky and E. Fuchs (2021). “Inflammatory adaptation in barrier tissues.” Cell 184(13): 3361–3375.

47. Okabe, J., C. Orlowski, A. Balcerczyk, C. Tikellis, M. C. Thomas, M. E. Cooper and A. El-Osta (2012). “Distinguishing hyperglycemic changes by Set7 in vascular endothelial cells.” Circ Res 110(8): 1067–1076.

48. Pandolfi, F., C. Brun-Buisson, D. Guillemot and L. Watier (2022). “One-year hospital readmission for recurrent sepsis: associated risk factors and impact on 1-year mortality-a French nationwide study.” Crit Care 26(1): 371.

49. Papavassiliou, A. G. and A. M. Musti (2020). “The Multifaceted Output of c-Jun Biological Activity: Focus at the Junction of CD8 T Cell Activation and Exhaustion.” Cells 9(11).

50. Peng, Y., Q. Wang, F. Jin, T. Tao and Q. Qin (2024). “Assessment of urine CCL2 as a potential diagnostic biomarker for acute kidney injury and septic acute kidney injury in intensive care unit patients.” Ren Fail 46(1): 2313171.

51. Pober, J. S. and W. C. Sessa (2007). “Evolving functions of endothelial cells in inflammation.” Nat Rev Immunol 7(10): 803–815.

52. Potter, D. R., J. Jiang and E. R. Damiano (2009). “The recovery time course of the endothelial cell glycocalyx in vivo and its implications in vitro.” Circ Res 104(11): 1318–1325.

53. Quinlan, A. R. and I. M. Hall (2010). “BEDTools: a flexible suite of utilities for comparing genomic features.” Bioinformatics 26(6): 841–842.

54. Renoux, F., M. Stellato, C. Haftmann, A. Vogetseder, R. Huang, A. Subramaniam, M. O. Becker, P. Blyszczuk, B. Becher, J. H. W. Distler, G. Kania, O. Boyman and O. Distler (2020). “The AP1 Transcription Factor Fosl2 Promotes Systemic Autoimmunity and Inflammation by Repressing Treg Development.” Cell Rep 31(13): 107826.

55. Riksen, N. P., S. Bekkering, W. J. M. Mulder and M. G. Netea (2023). “Trained immunity in atherosclerotic cardiovascular disease.” Nat Rev Cardiol 20(12): 799–811.

56. Saeed, S., J. Quintin, H. H. Kerstens, N. A. Rao, A. Aghajanirefah, F. Matarese, S. C. Cheng, J. Ratter, K. Berentsen, M. A. van der Ent, N. Sharifi, E. M. Janssen-Megens, M. Ter Huurne, A. Mandoli, T. van Schaik, A. Ng, F. Burden, K. Downes, M. Frontini, V. Kumar, E. J. Giamarellos-Bourboulis, W. H. Ouwehand, J. W. van der Meer, L. A. Joosten, C. Wijmenga, J. H. Martens, R. J. Xavier, C. Logie, M. G. Netea and H. G. Stunnenberg (2014). “Epigenetic programming of monocyte-to-macrophage differentiation and trained innate immunity.” Science 345(6204): 1251086.

58. Scarfe, L., D. Schock-Kusch, L. Ressel, J. Friedemann, Y. Shulhevich, P. Murray, B. Wilm and M. de Caestecker (2018). “Transdermal Measurement of Glomerular Filtration Rate in Mice.” J Vis Exp (140).

59. Schnitzler, J. G., R. M. Hoogeveen, L. Ali, K. H. M. Prange, F. Waissi, M. van Weeghel, J. C. Bachmann, M. Versloot, M. J. Borrelli, C. Yeang, D. P. V. De Kleijn, R. H. Houtkooper, M. L. Koschinsky, M. P. J. de Winther, A. K. Groen, J. L. Witztum, S. Tsimikas, E. S. G. Stroes and J. Kroon (2020). “Atherogenic Lipoprotein(a) Increases Vascular Glycolysis, Thereby Facilitating Inflammation and Leukocyte Extravasation.” Circ Res 126(10): 1346–1359.

60. Sessler, C. N., A. C. Windsor, M. Schwartz, L. Watson, B. J. Fisher, H. J. Sugerman and A. A. Fowler, 3rd (1995). “Circulating ICAM-1 is increased in septic shock.” Am J Respir Crit Care Med 151(5): 1420–1427.

61. Shankar-Hari, M., M. Ambler, V. Mahalingasivam, A. Jones, K. Rowan and G. D. Rubenfeld (2016). “Evidence for a causal link between sepsis and long-term mortality: a systematic review of epidemiologic studies.” Crit Care 20: 101.

62. Shankar-Hari, M. and G. D. Rubenfeld (2016). “Understanding Long-Term Outcomes Following Sepsis: Implications and Challenges.” Curr Infect Dis Rep 18(11): 37.

63. Shapiro, N. I., P. Schuetz, K. Yano, M. Sorasaki, S. M. Parikh, A. E. Jones, S. Trzeciak, L. Ngo and W. C. Aird (2010). “The association of endothelial cell signaling, severity of illness, and organ dysfunction in sepsis.” Crit Care 14(5): R182.

64. Sohrabi, Y., S. M. M. Lagache, V. C. Voges, D. Semo, G. Sonntag, I. Hanemann, F. Kahles, J. Waltenberger and H. M. Findeisen (2020). “OxLDL-mediated immunologic memory in endothelial cells.” J Mol Cell Cardiol 146: 121–132.

65. Sohrabi, Y., G. V. H. Sonntag, L. C. Braun, S. M. M. Lagache, M. Liebmann, L. Klotz, R. Godfrey, F. Kahles, J. Waltenberger and H. M. Findeisen (2020). “LXR Activation Induces a Proinflammatory Trained Innate Immunity-Phenotype in Human Monocytes.” Front Immunol 11: 353.

66. Song, D., Y. Lian and L. Zhang (2023). “The potential of activator protein 1 (AP-1) in cancer targeted therapy.” Front Immunol 14: 1224892.

67. Tavaci, T., Z. Halici, E. Cadirci, M. Ozkaraca and K. Kasali (2025). “The impact of tocilizumab treatment on the severity of inflammation and survival rates in sepsis is significantly influence by the timing of administration.” Inflammopharmacology 33(3): 1393–1405.

68. Venet, F. and G. Monneret (2018). “Advances in the understanding and treatment of sepsis-induced immunosuppression.” Nat Rev Nephrol 14(2): 121–137.

69. Vierbuchen, T., E. Ling, C. J. Cowley, C. H. Couch, X. Wang, D. A. Harmin, C. W. M. Roberts and M. E. Greenberg (2017). “AP-1 Transcription Factors and the BAF Complex Mediate Signal-Dependent Enhancer Selection.” Mol Cell 68(6): 1067–1082 e1012.

70. Wang, Q., M. Li, T. Wu, L. Zhan, L. Li, M. Chen, W. Xie, Z. Xie, E. Hu, S. Xu and G. Yu (2022). “Exploring Epigenomic Datasets by ChIPseeker.” Curr Protoc 2(10): e585.

71. Wang, T., A. Derhovanessian, S. De Cruz, J. A. Belperio, J. C. Deng and G. S. Hoo (2014). “Subsequent infections in survivors of sepsis: epidemiology and outcomes.” J Intensive Care Med 29(2): 87–95.

72. Weiss, E., A. Vlahos, B. Kim, S. Wijegunasekara, D. Shanmuganathan, T. Aitken, J. E. Joo, S. Imran, R. Shepherd, J. M. Craig, M. Green, U. Hiden, B. Novakovic and R. Saffery (2021). “Transcriptomic Remodelling of Fetal Endothelial Cells During Establishment of Inflammatory Memory.” Front Immunol 12: 757393.

73. Yan, F., D. R. Powell, D. J. Curtis and N. C. Wong (2020). “From reads to insight: a hitchhiker’s guide to ATAC-seq data analysis.” Genome Biol 21(1): 22.

74. Yu, H., L. Lin, Z. Zhang, H. Zhang and H. Hu (2020). “Targeting NF-kappaB pathway for the therapy of diseases: mechanism and clinical study.” Signal Transduct Target Ther 5(1): 209.

75. Yukawa, M., S. Jagannathan, S. Vallabh, A. V. Kartashov, X. Chen, M. T. Weirauch and A. Barski (2020). “AP-1 activity induced by co-stimulation is required for chromatin opening during T cell activation.” J Exp Med 217(1).

76. Zhang, H., Y. Wang, M. Qu, W. Li, D. Wu, J. P. Cata and C. Miao (2023). “Neutrophil, neutrophil extracellular traps and endothelial cell dysfunction in sepsis.” Clin Transl Med 13(1): e1170.

77. Zhang, Y., T. Liu, C. A. Meyer, J. Eeckhoute, D. S. Johnson, B. E. Bernstein, C. Nusbaum, R. M. Myers, M. Brown, W. Li and X. S. Liu (2008). “Model-based analysis of ChIP-Seq (MACS).” Genome Biol 9(9): R137.

78. Zhao, G. J., D. Li, Q. Zhao, J. X. Song, X. R. Chen, G. L. Hong, M. F. Li, B. Wu and Z. Q. Lu (2016). “Incidence, risk factors and impact on outcomes of secondary infection in patients with septic shock: an 8-year retrospective study.” Sci Rep 6: 38361.

79. Zhou, G., J. Liu, H. Zhang, X. Wang and D. Liu (2022). “Elevated endothelial dysfunction-related biomarker levels indicate the severity and predict sepsis incidence.” Sci Rep 12(1): 21935.

