## Supplementa Figures for "Endothelial Cells retain inflammatory memory through chromatin remodeling"

### Supplemental Figures

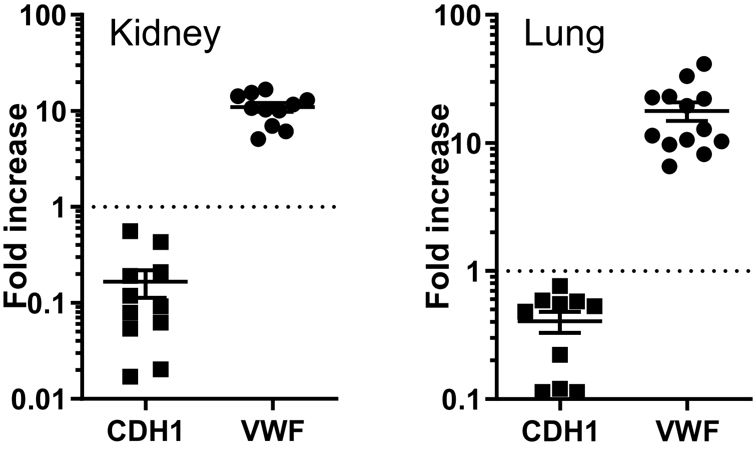


**Supplemental Fig 1.** Validation of EC enrichment from kidney and lung.
RT-qPCR analysis of sorted ECs from kidney and lung demonstrates enrichment of endothelial marker vWF and depletion of epithelial marker CDH1. Data shown as fold change calculate by the endothelial enrichment over the total RNA. Mean ± SEM; log scale.

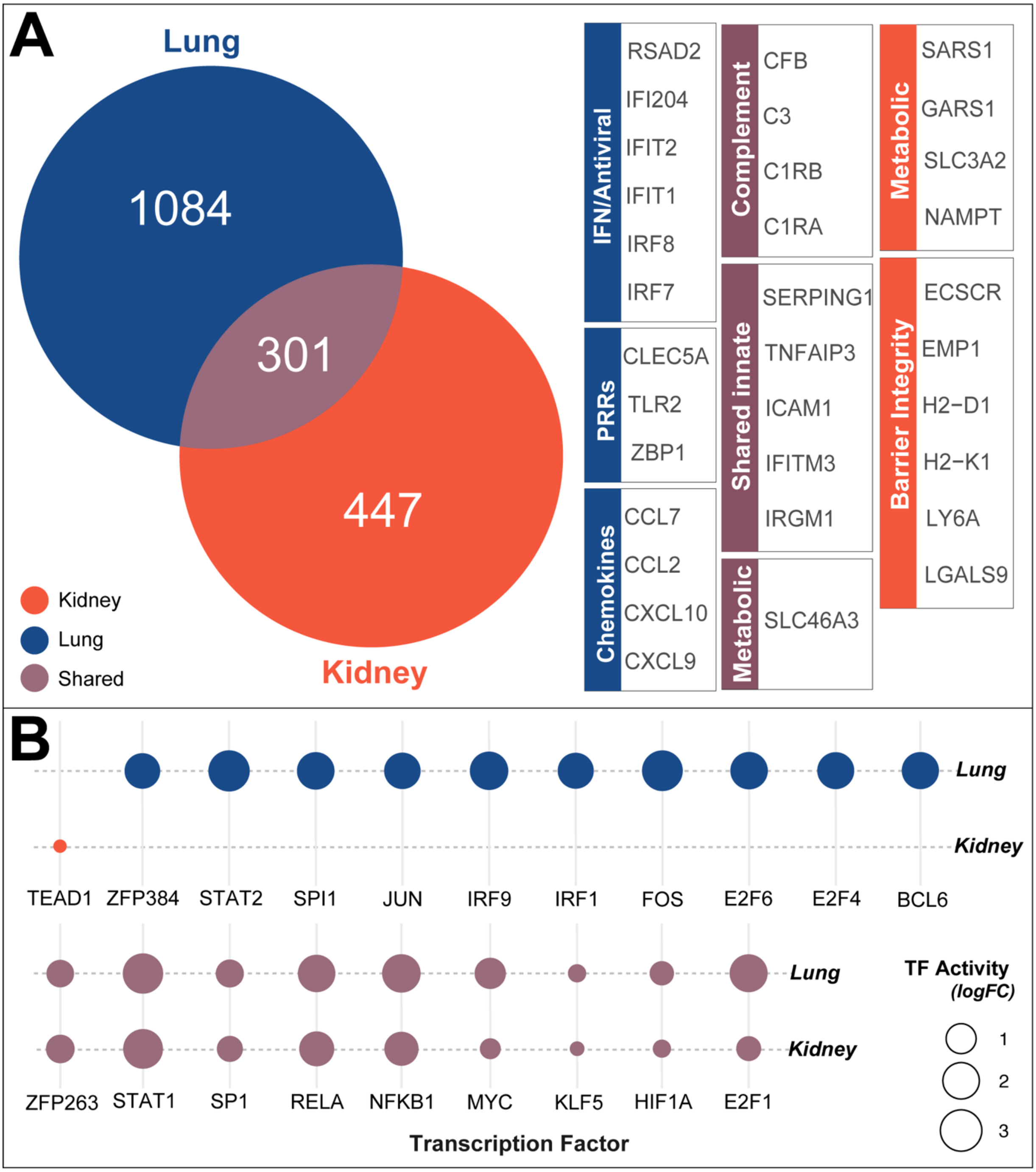


**Supplemental Fig 2.** Supplemental Figure 2. Shared and organ-specific transcriptional memory genes.

**(A)** Venn diagram and heatmap classification of 301 shared and 1084 lung- or 447 kidney-specific genes amplified in CLP+SP ECs. **(B)** Bubble plot of transcription factor (TF) activity predicted by DoRothEA analysis in shared and organ-specific gene sets. Size of each bubble represents TF activity score; color denotes tissue origin.

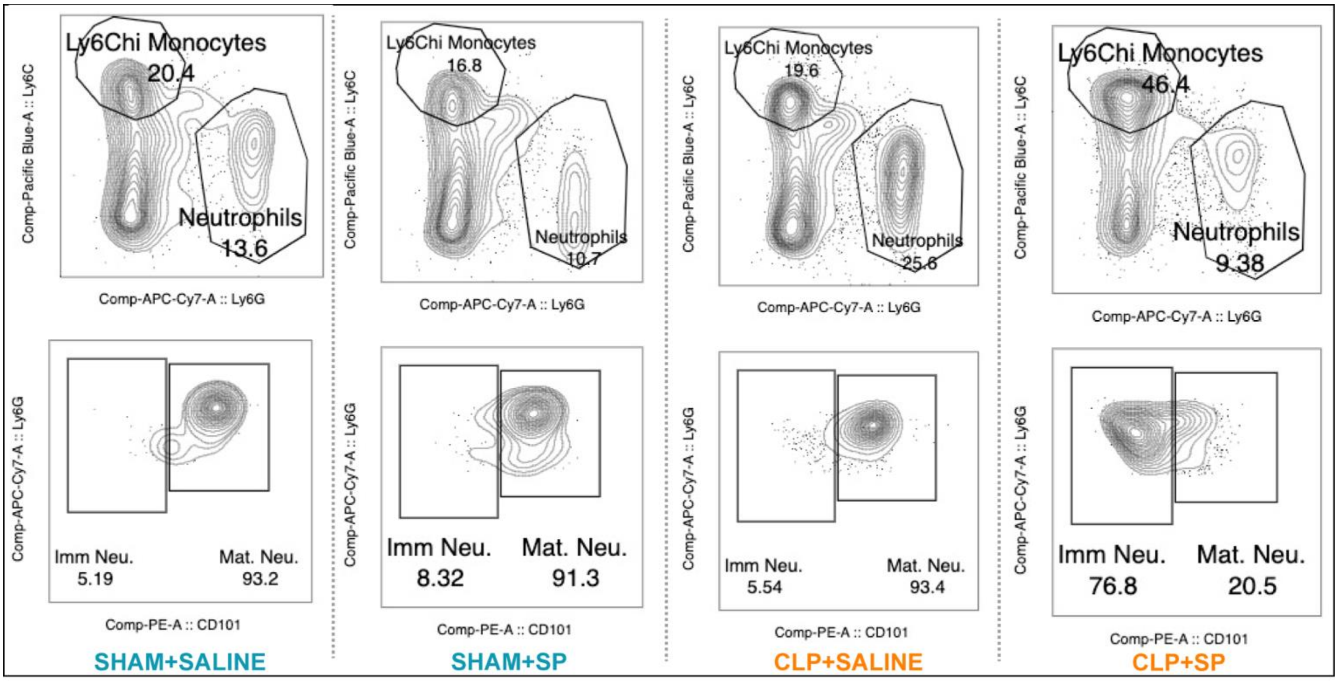


**Supplemental Fig 3.** Representative flow plots for CD45$^{+}$Ly6G$^{+}$ neutrophils, CD11b$^{+}$Ly6C^hi monocytes, and CD101$^{-}$ immature neutrophils across all groups. These define the strategy for quantifying myeloid cell subsets in Figure 3.

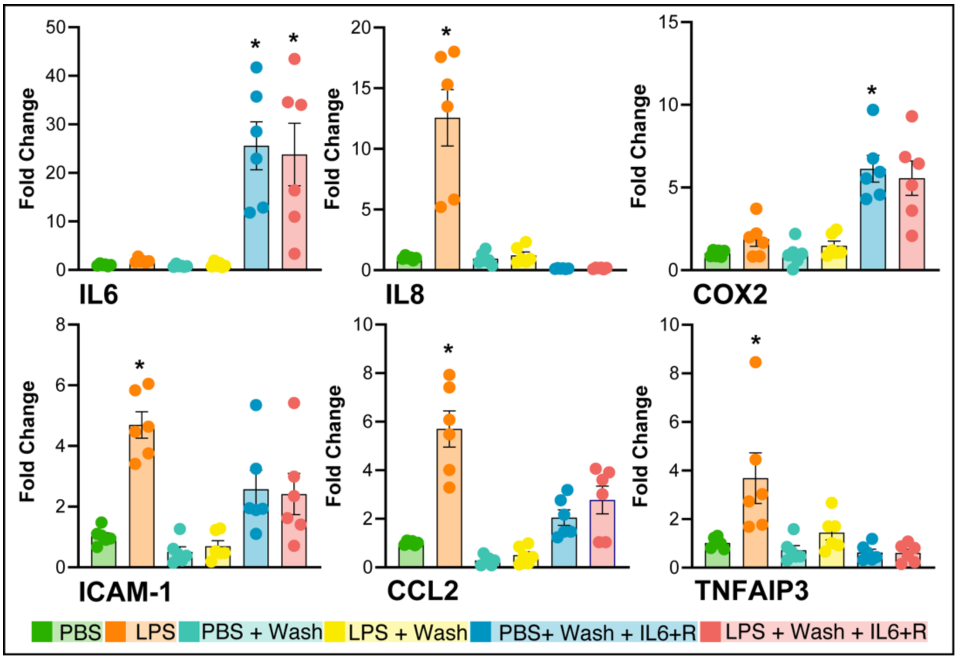


**Supplemental Fig 4.** Inverted two-hit model reveals absence of amplified inflammatory response when IL-6 is delivered after LPS.
HUVECs were treated with LPS (1 µg/mL) or PBS for 72 hours (1st hit), followed by a 48-hour wash and subsequent 6-hour exposure to IL-6+R (200 ng/mL) or PBS (2nd hit). Gene expression was measured by qPCR and is presented as fold change relative to PBS control. Data are mean ± SEM; *p<0.05 vs. PBS control, one-way ANOVA with Tukey's post hoc test.
